# Long-distance gene flow and contrasting population structures of reef-building corals and their algal symbionts inform adaptive potential across the Western Pacific

**DOI:** 10.1101/2025.10.14.682263

**Authors:** Hugo Denis, Katharine E. Prata, Hisatake Ishida, Iva Popovic, Véronique J.L. Mocellin, Magali Boussion, Ilha Byrne, Steven W. Purcell, Line K. Bay, Gaël Lecellier, Cheong Xin Chan, Cynthia Riginos, Emily J. Howells, Véronique Berteaux-Lecellier

## Abstract

The genetic diversity and connectivity of reef-building coral populations are key to their survival in warming oceans. Yet our understanding of corals’ demographic resilience and adaptive potential is complicated by cryptic species diversity, wide geographic distributions, and complex coral-algal symbioses. To address these challenges, we investigated genetic connectivity and diversity of the broadcast-spawning coral *Acropora spathulata* and its associated Symbiodiniaceae across 29 reefs spanning the Great Barrier Reef, the Coral Sea, and New Caledonia, using whole-genome sequencing of 1,088 colonies. We identified four genetically distinct coral populations that diverged between 0.27 and 0.65 million years ago, likely due to geographic isolation across thousands of kilometers. These populations maintained asymmetrical gene flow along major ocean currents despite demographic isolation, and sustained large local effective population sizes (∼2,900), supported by a high dispersal range of ∼100 km per generation. In contrast, their Symbiodiniaceae partners varied over finer spatial scales, with five distinct *Cladocopium* taxa distributed along latitudinal and cross-shore gradients, likely driven by local environmental conditions. These results suggest that high dispersal capacity and large local population size promote demographic resilience within reef systems, while environment-specific symbioses and long-distance gene flow across reef-systems support adaptation and evolutionary rescue.

## Introduction

Reef-building corals and their microbial symbionts constitute coral holobionts that are the foundation of biodiverse but threatened coral reef ecosystems (Souter et al., 2021). As tropical corals are declining globally, it is crucial to better understand the evolutionary, spatial and environmental processes that shape their population dynamics and resilience (Edmunds & Riegl, 2020; van Oppen & Gates, 2006). However, the widespread presence of cryptic coral diversity (Grupstra et al., 2024; Riginos et al., 2024), along with vast geographic ranges (Gleason & Hofmann, 2011), challenges our ability to determine the size and connectivity of coral populations. In addition, variability in symbioses between coral hosts and symbiotic microalgae from the family Symbiodiniaceae can affect holobionts’ capacity to grow, survive and adapt under changing conditions (Bourne et al., 2016).

Population genomics, which investigates genome-wide molecular variation, can differentiate coral taxa that are indistinguishable based on morphological criteria (Bongaerts et al., 2021), or traditional population genetic markers such as microsatellites (Arrigoni et al., 2020; Meziere et al., 2024). Within coral species, it can provide a holistic view of spatial genetic variation patterns in both hosts and symbionts across their range (Fuentes-Pardo & Ruzzante, 2017). These analyses are essential to uncover the distinct scales at which gene flow, local adaptation and co-evolutionary dynamics influence holobionts adaptive responses.

Contemporary gene flow (over tens of generations) is crucial for coral demographic dynamics, particularly for the recovery of locally depleted coral populations following perturbations (Hock et al., 2017; Holbrook et al., 2018). To quantify its spatial extent, isolation-by-distance (IbD) theory provides a useful framework by estimating a generational gene dispersal distance (Rousset, 1997; Wright, 1946). Combining the slope of IbD with estimates of effective density, one can infer the generational mean axial dispersal distance (σ), over which most dispersal events occur, and vast differences in σ depending on reproductive modes and taxa have been described (Gorospe & Karl, 2013; Japaud et al., 2019; Meziere et al., 2025; Prata et al., 2024). To estimate effective density, the local effective population size (*N_e_*) is estimated and divided by the area the population occupies. Local *N_e_* indicates the loss rate of genetic variation due to genetic drift at the scale where most breeding events occur (Waples, 2024). This standing variation on which natural selection acts, underpins adaptive potential (Fisher, 1930). Jointly estimating σ and *N_e_* thus provides crucial information on the extinction risk of local coral populations (Hernández-Agreda et al., 2024) and the spatial scale of demographic replenishment (Gaines et al., 2010), both are essential for defining conservation units.

Over longer evolutionary time-scales, occasional long-distance dispersal can affect species range and adaptation to global changes (Clobert, 2012). However, methods based on high-level genetic differentiation cannot disentangle the effect of migration from genetic drift and shared ancestry, or determine direction of gene flow (Lawson et al., 2018; Whitlock & McCauley, 1999). Demographic history inference approaches that model divergence times, variation in migration rates and long-term effective population sizes (Gutenkunst et al., 2009) offer a complimentary strategy to assess gene flow across a coral species range (Meziere et al., 2025). However, despite potential for coral connectivity spanning thousands of kilometers (Davies et al., 2015; Matz et al., 2018), such methods have been mostly applied within national borders (Matz et al., 2018; Meziere et al., 2025; Tsuchiya et al., 2022; Zhang et al., 2022). This limits the development of coordinated regional conservation strategies that incorporate gene flow from different regions to assess evolutionary adaptation potential of corals (Colton et al., 2022).

Importantly, coral evolution and conservation require consideration of their symbiosis with Symbiodiniaceae microalgae (LaJeunesse et al., 2018) that are genetically and physiologically diverse (Nitschke et al., 2022). For instance, photosynthetic ability of Symbiodiniacae is known to differ among taxa under heat stress (Levin et al., 2016); some taxa inhabit warmer environments (McRae et al., 2023) or increase in prevalence during thermal stress (LaJeunesse et al., 2009). Most Symbiodiniaceae are facultative symbionts, therefore both macro-environments and host micro-environments will favor certain taxa or genotypes (Bhattacharya et al., 2024; Quigley et al., 2019; Reich et al., 2021). In turn, some symbiont taxa may facilitate corals’ local adaptation to environmental conditions (Howells et al., 2012). Symbiont diversity has been mostly described using phylogenetic markers such as the ribosomal internal transcribed spacer 2 (ITS2; Hume et al., 2019) and the chloroplastic non-coding region of *psbA* gene, i.e., psbA^ncr^ (Moore et al., 2003), or via multilocus genotyping (e.g., Davies et al., 2020; Lewis et al., 2024). Yet, distinguishing intra-from inter-genomic variants remains challenging, and the dependence on few selected marker genes may limit taxonomic resolution (Armstrong et al., 2024). Methods based on genome-wide sequence composition of hologenome samples are thus emerging (Ishida et al., 2024), unlocking hidden phylogenetic (Armstrong et al., 2024; Cooke et al., 2020; Zhang et al., 2022) and ecological (Hoadley et al., 2021) diversity of these coral symbionts.

*Acropora spathulata* is a fast-growing, broadcast-spawning and widespread species that inhabits shallow reef environments in the Western Pacific, including the Great Barrier Reef (GBR), the Coral Sea, and New Caledonia (NC). Genomic data has revealed new insights into the cryptic diversity of *Acropora* corals on the GBR (Matias et al., 2022; Naugle et al., 2024), their associated Symbiodiniaceae (LaJeunesse et al., 2004; Matias et al., 2022; Naugle et al., 2024; van Oppen et al., 2001) and realized connectivity, dominated by southward migration of coral larvae (Matz et al., 2018; Riginos et al., 2019). However, such information remains scarce in NC with only one genomic study conducted to date on *Acropora* coral hosts (Selmoni et al., 2021), and few on their symbiont communities (Alessi et al., 2024; Camp et al., 2020; Denis et al., 2024b). In addition, despite being one of the most studied coral reef ecosystem (Alvarado-Cerón et al., 2022), GBR’s connectivity with nearby western Pacific islands such as NC remains unknown. Several ocean currents flow westward and eastward between the GBR and NC (Figure S1; Cravatte et al., 2015), offering potential for periodical dispersal via stepping-stone migration through Coral Sea atolls, e.g. the Chesterfield-Bellona (CB) plateau. To our knowledge, no study—in corals or other marine organisms—has systematically assessed gene flow in this western Pacific region, aside from pumice rafting coral larvae from NC to the GBR (Jokiel, 1990), and some genetic admixture between these reef systems and Coral Sea atolls (Lukoschek et al., 2016; Oury et al., 2020). NC and the Southern GBR are often considered potential climate refugia with projected stability of coral cover (Marzonie et al., 2025; Matz et al., 2020; Sully et al., 2022) due to the mild occurrence of bleaching until recently (Shlesinger & van Woesik, 2023). Understanding whether these systems have or still exchange migrants is therefore important for their regional conservation.

Here, we systematically assess the host and symbiont population structures and gene flow of the coral *Acropora spathulata*, expanding the common research scope of the GBR (Cooke et al., 2020; Matias et al., 2022; Scott et al., 2024; van Oppen et al., 2018; Warner et al., 2015) to the Western Pacific. By integrating existing and innovative genome-scale approaches, we densely sampled, sequenced, and analyzed whole-genome sequencing data from 1,088 colonies from 29 reefs across the GBR, the Coral Sea and NC. Our study represents one of the most geographically comprehensive efforts to concurrently investigate coral host and symbiont diversity as well as contemporary and long-term connectivity.

## Methods

### Study sites and sample collection

*A. spathulata* was sampled from the east coast of Australia to the west coast of New Caledonia and across a 10 ° latitudinal gradient (Table S1). The first batch of samples consisted of 831 colonies collected across 15 reefs of the Great Barrier Reef (*n* = 40–60 per reef; Figure 1) between 28 February and 26 March 2022 (GBRMPA permit G21/45166.1) as described in Denis et al. (2024a). The second batch comprised 323 colonies collected across 14 reefs of the western coast of New Caledonia (NC) and atolls of the Chesterfield-Bellona (CB) plateau located in the Coral Sea (*n* = 4–25 per reef) between 26 June and 28 September 2023 (New Caledonia Natural Park of the Coral Sea permit 2023-1503/GNC, South Province permit 4508-2022/ARR/DDDT and North Province permit 609011). At each reef, small fragments (< 1 cm^3^) were collected on SCUBA from coral colonies originating from one or two sites (covering 50–4200 m^2^) and immediately stored in 80–100% ethanol for DNA extraction. Depending on reef sites geomorphology, colonies were collected from upper reef slopes, fringing reefs, inner barrier reefs or reef flats at an average depth of 0.6 m (-0.7 m to 5.6 m relative to the lowest astronomical tide). Geographic coordinates were recorded for each colony by carrying a GPS in tow on a surface float (except at NC and CB reefs), and whole colony photographs and macrophotographs were taken of each colony using an Olympus TG-6. The occurrence of clones was minimized by avoiding sampling neighboring colonies of similar pigmentation.

### DNA extraction and sequencing

Coral fragments from 1,154 colonies were used for DNA extraction and 1,144 high quality DNA samples were used for library preparation and whole genome sequencing (Table S2). While extraction and sequencing of the GBR and New Caledonia samples were conducted at different facilities, similar extraction and library preparation kits and sequencing platforms were used in order to minimize batch effects. In addition, a subset of 23 samples from the GBR were extracted and sequenced alongside both batches to control for such effects (Supplementary Material and Methods). In both cases, total DNA was extracted from coral tissue slurry (tissue and skeleton) using Qiagen DNeasy Blood and Tissue Kit (Qiagen, Hilden, Germany) following manufacturer’s instruction, including an RNAse digestion step for GBR samples (30 min at room temperature). For GBR samples, sequencing libraries were prepared using the Lotus DNA Library Prep Kit for NGS (10 ng of input DNA, 8 PCR cycles) and sequenced on the Illumina NovaSeq 6000 platform (200 cycles; 150 bp paired end reads; targeting 10X coverage; Azenta Life Sciences, CSIRO). For New Caledonia samples, the libraries were prepared using the Truseq Nano DNA kit (200 ng of input DNA, 8 PCR cycles) and sequenced on the Illumina NovaSeq X platform (300 cycles; 150 bp paired end reads; targeting 10X coverage; Macrogen, Korea).

**Figure 1.**
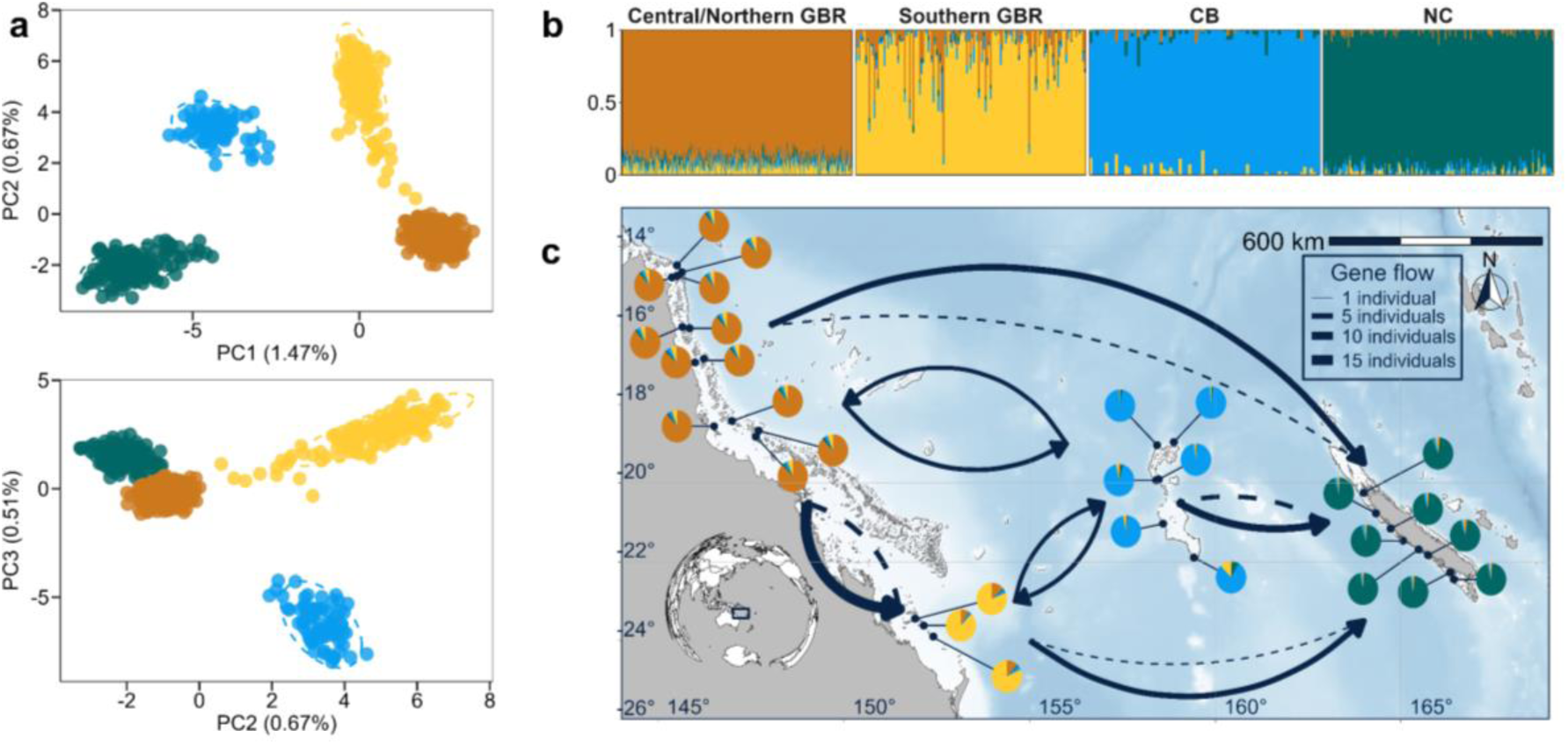
Population genomic structure and connectivity of *Acropora spathulata* across reefs from the Great Barrier Reef, Chesterfields-Bellona atolls (CB) and New Caledonia (NC). (a) Principal component analysis based on 27,175 SNPs showing four genetically and geographically distinct populations across the western Pacific. PC1, PC2 and PC3 are shown with the proportion of variance explained by each component indicated in percentages. (b) Ancestry proportions from each individual using ADMIXTURE analysis with the optimal number of ancestral populations found using cross-validation (K=4). (c) Map showing ancestry proportion at sampling sites, averaged across colonies (n=4–60). Arrows indicate average gene flow, in number of migrants per generation, between populations since their divergence inferred from demographic modeling in dadi. Solid and dashed lines indicate the primary and secondary directions of gene flow, respectively.

### Genomic data pre-variant processing and filtering

The bioinformatic pipeline used to process, filter, and analyze the genomic data is outlined in Figure S2. The dataset consisted of the 1,144 sequenced individuals with technical replicates (*n* = 25; Supplementary Material and Methods). Pre-variant filtering steps are summarized in Table S2. Adapter removal and read quality trimming was conducted using *Trimmomatic* v0.39 (Bolger et al., 2014) with a 4 bp sliding window, a minimum phred score quality of 20, and a minimum read length of 50 bp. Leading and trailing low quality bases were removed (LEADING:3, TRAILING:3) and adapter sequences were removed using the Illuminaclip option in ‘palindrome mode’ (2:30:10:4). Sample quality was confirmed using *FastQC* v0.12.1 (http://www.bioinformatics.bbsrc.ac.uk/projects/fastqc) and *MultiQC* v1.21 (Ewels et al., 2016). Filtered reads were then aligned to the *Acropora millepora* reference genome v3 assembly (unpublished upgrade from GCA_013753865.1_Amil_v2.1; see Supplementary Material and Methods for a justification of the reference genome choice) using the *Burrow-Wheeler Aligner* v0.7.17 (Li & Durbin, 2010) and the *MEM* algorithm at default settings. SAM files were converted to indexed and sorted BAM files using *Samtools* v1.20 (Danecek et al., 2021) and PCR duplicates were marked and removed using *Picard* v3.1.1 (VALIDATION_STRINGENCY=LENIENT; http://broadinstitute.github.io/picard/). After filtering of samples that failed to be sequenced and samples with unexpectedly low percentage (< 80%; possibly indicative of mis-identified taxa) or number (< 10M) of mapped reads, 1,132 BAM files were retained. To provide an initial evaluation of population structure across the study system and pinpoint potential clones and taxa misidentification, this dataset was analyzed with a genotype likelihood framework in *ANGSD* v0.940 (Korneliussen et al., 2014; see Supplementary Material and Methods for results). Four samples from the GBR and 23 samples from NC were discarded as they likely belonged to *A. millepora* and a distinct unknown taxon, respectively.

### SNPs calling, clone identification and filtering

Indexed, duplicate-free BAM files were used to call variants and genotypes from a total of 1,105 remaining samples using *GATK* v4.5.0.0 ‘Germline short variant discovery best practice’ workflow (Van der Auwera & O’Connor, 2020). Variant calling was first performed per sample, focusing only on reads mapping to the 14 reference chromosomes, using the *HaplotypeCaller* tool that performs *de novo* assembly of haplotypes in active regions. Individual ‘gvcf’ files were then consolidated per chromosome using *GenomicDBimport* and used for joint genotyping using *GenotypeGVCFs* tool. *GatherVCFs* and *SelectVariants* tools were used to combine chromosome files in a single variant call format (VCF) file and separate SNPs from INDELs and monomorphic sites. The SNPs VCF file was preliminarily filtered using *GATK VariantFiltration* tool following GATK stringent filtering best practices (QD > 2, FS < 50, MQ > 40, MQRankSum > -12.5, ReadPosRankSum > -8, SOR < 3, GQ > 20) and 17 individuals with high missingness (> 85%) were discarded. Previously identified putative clonal colonies were confirmed and filtered to retain only one colony with the lowest missingness per clonal group (Supplementary Material and Methods, Table S3).

### Host population genomics analyses

To confirm the absence of strong batch effects that could confound population structure, a subset of 23 technical replicates pairs was sequenced along both GBR and NC samples (Supplementary Material and Methods; Figure S3). After exclusion of natural clones and technical replicates, a final set of 999 individuals was used for host population genetic analyses. A first dataset of unlinked SNPs was created to investigate population structure and phylogenetic relationship between populations of the Coral Sea, retaining only bi-allelic SNPs with a minimum mean locus depth of 5 and a maximum of 15, a minor allele frequency (MAF) > 0.05 and a locus missingness < 20% with *vcftools* v0.1.17 (Danecek et al., 2011). In addition, SNPs in linkage disequilibrium with a pairwise *r^2^* > 0.2 were removed using *plink* v1.90b7.2 (Purcell et al., 2007; “Dataset 1”, Table S4) because we were interested in identifying neutral population structure patterns that occur throughout the genome. To identify genetically distinct groups of individuals, we performed a principal component analysis (PCA) with the *glPca* function from R package *adegenet* v2.1.10 (Jombart, 2008). We then estimated individual ancestry proportion using ADMIXTURE v1.3.0 (Alexander & Lange, 2011). The optimal number of ancestral populations (*K*=4) was determined using the cross-validation method implemented in ADMIXTURE. Following Hemstrom et al. (2024) best practices, these analyses were also conducted using variable stringency MAF (MAF > 0.01 and MAF > 0.05) and missingness thresholds (5%, 10%, 20% and 50%).

Finally, we used a subset of individuals with the highest ancestral assignment in ADMIXTURE (200 colonies, 50 colonies per population) and built an individual-based Maximum Likelihood phylogenetic tree using RAxML (Stamatakis, 2014). To build the tree, the VCF file was converted to the Phylip format using *vcf2phylip.py* script (https://github.com/edgardomortiz/vcf2phylip) and raxmlHPC-PTHREADS was run with a GTRGAMMA model and 100 bootstrap replicates.

### Population statistics

A second dataset of linked SNPs was used to compute population genetic differentiation and diversity estimates retaining SNPs with a minimum mean locus depth of 5 and a maximum of 50, a minimum allele count (MAC) of 1, a *p*-value of deviation from HWE test >0.0001 and < 20% missingness (“Dataset 2”, Table S4) following recommendations of Hemstrom et al. (2024). To generate unbiased estimates of population genomic statistics despite missing data, the SNP dataset was concatenated to include invariant sites, filtered with the same depth and missingness thresholds per above, and used to compute statistics with *pixy* v1.2.10 (Korunes & Samuk, 2021). Following Sopniewski and Catullo (2024), 10 replicates of 5 randomly selected individuals per population were used to compute the fixation index (*F_st_*) using the Weir and Cockerham method (Weir & Cockerham, 1984). Because *F_st_* estimates can be affected by within population diversity, we also computed absolute divergence (*D_xy_*) for each replicate, to get more-independent estimates of population differentiation. Finally, we computed nucleotide diversity (π) within each population. We reported the median *F_st_*, *D_xy_* and π estimates across replicates.

### Demographic modeling

To investigate the evolutionary history and connectivity of these populations, a subset of Dataset 2 comprising 120 colonies (30 colonies per population, showing >95% admixture cluster assignment) was used to conduct demographic modeling analyses in *dadi* v2.3.3 (Gutenkunst et al., 2009). This dataset was further filtered to retain SNPs with a minimum allele count (MAC) of 3 to minimize the effect of sequencing errors. SNPs showing a deviation from HWE (*p*-value < 0.0001) were also removed as we observed an excess in heterozygosity creating a peak in the SFS at 0.5 haplotype frequency, potentially due to gene duplication, paralogous loci or balancing selection. Finally, a second round of filtration to ensure < 20% missingness within each population was performed using *vcf_minrep_filter.py* script (https://github.com/pimbongaerts/radseq, “Dataset 3”, Table S4). VCF files were created for each population pair (n=6) and analyzed using a custom workflow to estimate effective population sizes, divergence time and migration rates between populations (https://github.com/kepra3/kp_dadi).

We computed joint-allelic frequency spectrums (JAFS) for each pair using the subsample method to account for missing data, and masking singletons and doubletons to minimize the effect of sequencing errors or somatic mutations. We initially tested three scenarios; divergence in isolation, divergence with asymmetrical migration and ancestral asymmetrical migration. We optimized the models’ parameters to best fit the observed JAFS (Figure S4). To avoid getting stuck in local optima, optimization was performed by exploring the parameter space in three stages: first with three-fold, then two-fold, and finally one-fold perturbations of the starting parameters, based on the previously optimized peak values (>100 runs per stage). The final set of parameters was chosen when appearing several times with 1-fold perturbation from different initial values. To get confidence intervals for these parameters, 100 bootstraps were created for each pair by resampling genomic contigs. Optimization runs (n=50) were performed for each bootstrap and model, using 1-fold perturbations starting from previously optimized parameters. For each bootstrap replicate, delta-AIC values were computed between models, and the mean delta-AIC across replicates was used to identify the best-fitting model for each population pair (Figure S5). Optimized parameters from the best-fitting model (divergence with asymmetrical migration) were used to calculate 95% credible intervals, after excluding bootstrap replicates within the lowest 10% likelihood quantile.

Finally, we converted parameter values obtained in *dadi* (𝜃, *v*_1*dadi*_, *v*_2*dadi*_, *M*_12*dadi*_, *M*_12*dadi*_ and *T*_*dadi*_) into physical units: divergence time in years, effective population sizes in number of individuals and gene flow between taxa in number of migrants per generation. To do so we first calculated the effective sequence length for each pair as:

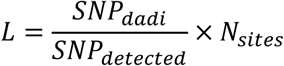

where 𝑆𝑁𝑃_*dadi*_ is the number of SNPs used to construct the JAFS from the specific pair, 𝑆𝑁𝑃_*d*𝑒𝑡𝑒𝑐𝑡𝑒*d*_ is the number of SNPs originally detected before filtration, and 𝑁_𝑠*i*𝑡𝑒𝑠_ is the number of sites obtained from GATK pipeline (including SNPs, INDELs and monomorphic sites). Next, we computed the ancestral population size (𝑁_𝑟𝑒𝑓_) and the effective population sizes *v*_1_ and *v*_2_ in number of individuals, the divergence time (*T*) in years, migration rates in fraction of migrant individuals at each generation (𝑚_12_; from Population 2 to Population 1 and 𝑚_21_; from Population 1 to Population 2) and rates of gene flow in number of individuals per generation (*M*_12_ and *M*_21_), using the following formulas:

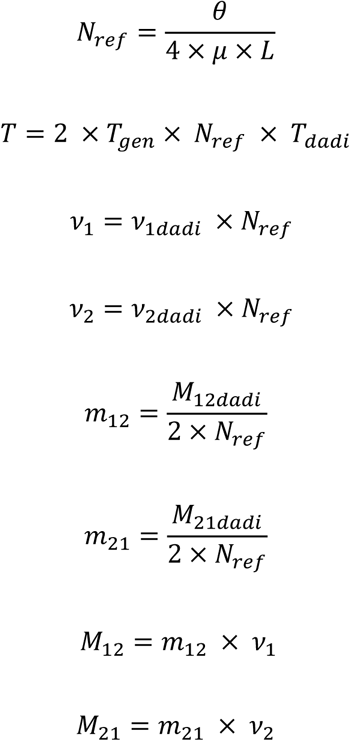

We used a mutation rate 𝜇 of 1.2 × 10^-8^ mutations per base per generation, previously estimated for *Acropora* corals (Zhang et al., 2022) and a generation time *T*_𝑔𝑒𝑛_ of 5 years following the assumptions of Matz et al. (2018) on generation times of *A. millepora*.

### Isolation by distance analysis and estimating N_e_

To estimate realized dispersal and effective population sizes, Dataset 2 was further filtered to retain SNPs with <10% missing data and physically pruned to remove SNPs separated by <500 bp (“Dataset 4”). This was done because both isolation by distance (IbD) analyses and *N_e_* estimation methods require no physical linkage (Waples, 2024).

Using this dataset, we estimated σ that represents ecologically relevant gene flow averaged across the last 10–15 generations, following Prata et al. (2024) with modifications. Multiple IbD regressions were performed in *spagedi* v1.5 (Hardy & Vekemans, 2002) using the empirical estimate of Rousset’s individual genetic distance (â; Rousset, 1999) and the geographic distance between individuals.

Geographic distances were calculated using the Haversine formula, that estimates great-circle distances between latitude-longitude pairs. Regression analysis conducted within each of the four populations (corresponding to geographic regions; Figure 1) revealed no significant IbD. Because we were interested in a general dispersal estimate across the species range and the *F_st_* estimates amongst populations were low, we used the global dataset for conducting the IbD analysis. Different regressions were performed to estimate the slope β for a range of spatial scales that roughly evenly partitioned the dataset (maximum distance 50 m to 2,300 km). We queried multiple spatial scales in order to select *a posteriori* the appropriate regression distance spanning minimum σ and maximum 10–50 σ following Hardy and Vekemans (1999). The dimensionality of the habitat was determined for each regression by the width to area ratio, using a 1D analysis for a ratio <0.1 and a 2D analysis and the log_10_ of geographic distance for a ratio >0.1 (Rousset, 1997). Similarly, for multiple spatial scales, the population density (D_e_) was estimated by dividing the effective population size by the habitat area (*A*) for 2D or length (*L*) for 1D regressions (see below). Confidence intervals for β were estimated using a jackknifing procedure across individuals and the slope was deemed significant when the confidence intervals did not span 0. All β and D_e_ estimates were then combined to estimate σ at each spatial scale using the following isolation-by-distance model equation (Rousset, 1997; Wright, 1946), example for two-dimensions:

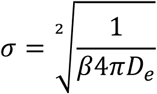

The linkage disequilibrium method was used to estimate the effective population size (*N_e_*) at the neighborhood extent (radius of 2𝜎), which *N_e_* represents the effective number of individuals that can easily mate in a continuously distributed population (*N_b_*; Neel et al., 2013). Neighborhood *N_e_* was estimated using *LDNe* v2.1 (Waples & Do, 2008) with a minimum allele frequency filter based on the number of samples following authors recommendations. Confidence intervals were obtained for *N_e_* using the jackknifing procedure.

Finally, we filtered our β, σ and *N_e_* estimates to ensure that the spatial scales at which the IbD regression was performed matched a σ value ranging from σ to 10–50σ following Hardy and Vekemans (1999). We also removed any instance where *N_e_* was not estimated at the neighborhood area, *A* or *L* equating to 4𝜋σ^2^ for two dimensional habitat (or 2√𝜋𝜎^2^ for one dimension) because this is the ideal spatial scale to estimate *N_e_* (Neel et al., 2013). Upper and lower confidence intervals for filtered estimates were then used to obtain distributions of β and *N_e_* through resampling (*n*=100). A gamma distribution was used for *N_e_* and a log-normal distribution for β. These distributions were then combined to create a joint 𝜎 distribution and propagate the error from each separate analysis.

### Symbiodiniaceae community analyses

We used genomic reads from 1,088 *A. spathulata* holobionts to characterize symbiont community composition and diversity across our study area, adopting the approach by Ishida et al. (2024), available at https://github.com/hisatakeishida/Symb-SHIN. Genomic reads not mapping to the host reference genome (i.e. non-coral reads) were first re-aligned to several reference genomes of Symbiodiniaceae using the Burrow-Wheeler Aligner v0.7.17(Li & Durbin, 2010) and the MEM algorithm (*bwa-mem*) at default setting, with one reference per genus (*Symbiodinium microadriaticum* CCMP2467 (Nand et al., 2021), *Breviolum minutum* Mf1.05b (Shoguchi et al., 2013), *Cladocopium proliferum* SCF055-01 (Chen et al., 2022), *Durusdinium trenchii* CCMP2556 (Dougan et al., 2024), *Effrenium voratum* RCC1521 (Shah et al., 2024) and *Fugacium kawagutii* CCMP2468 (Li et al., 2020)).

For the first-pass analysis of the symbiont community composition, we retrieved and classified ITS2 sequences present in the samples using *graftM* (Boyd et al., 2018). All ITS2 sequence variants were classified as belonging to genus *Cladocopium*. Consequently, to improve intra-specific taxonomic resolution we also analysed *psbA*^ncr^ sequences among the samples. We retrieved *psbA*^ncr^ sequences by mapping reads to a custom *psbA*^ncr^ reference database using *bwa-mem*. We then follow Armstrong et al. (2023) to retain only reads that aligned in full length (100% cover) at >90% identity, and used number of uniquely mapped reads as proxy for abundance.

To capture genetic diversity of symbionts directly using whole-genome sequencing data not restricted to selected marker genes, we adopted a *k*-mer-based alignment-free approach, following earlier studies (Ishida et al., 2024; Zhang et al., 2022). The *k*-mers in each sample were first enumerated using *jellyfish* v1.1.12 (https://github.com/gmarcais/Jellyfish) at *k* = 21 yielding a 21-mer profile following previous studies (González-Pech et al., 2021). Pairwise distance between samples was computed from these profiles using *d2ssect* (https://github.com/bakeronit/d2ssect), based on D ^s^ metric (Reinert et al., 2009). Samples containing similar symbiont compositions are expected to share similar *k*-mer profiles (i.e., short D ^s^-derived distance).

To validate the aforementioned methods using a conventional amplicon-based approach, a subset of 36 DNA samples distributed across host populations (Figure S6) were used for ITS2 marker amplification (Herculase II Fusion DNA Polymerase Nextera XT Index V2 Kit) and Illumina sequencing at Macrogen (Seoul, Korea) using forward primer SYM_VAR_5.8S 5’-GAATTGCAGAACTCCGTGAACC-3’ and reverse primer SYM_VAR_REV 5’-CGGGTT CWCTTGTYTGACTTCATGC-3′ (Hume et al., 2018). ITS2 sequence data was submitted to the SymPortal analytical framework (https://symportal.org; Hume et al., 2019) and ITS2 sequence variants and type profiles output by the pipeline were used to visualize the results. For this subset of 36 individuals, *graftM* results were checked for consistency with SymPortal results using a relative abundance barplot of ITS2 sequence variants. Using the same set of samples, we also compared the hierarchical clustering of Unifrac distances based on ITS2 ‘defining intragenomic variants’ with that based on D ^s^ distances, which showed great consistency (Figure S7) and thus we assigned each of the D ^s^ clusters the corresponding major ITS2 sequence. Both dendrograms were aligned to build a tanglegram with R package *dendextend* and the stepside method (Galili et al., 2015).

### Host and symbiont co-clustering analysis

To visualize how Symbiodiniaceae communities varied across host populations, we performed a hierarchical clustering of samples using the pairwise distance matrix derived from the D ^s^ metric (above) using all samples (*n* = 878) for which > 50 M bp of reads mapped to the reference Symbiodiniaceae genomes. This dendrogram was pruned to build a tanglegram with the host phylogenetic tree obtained in *RAxML* using *dendextend*.

### Distance based redundancy analysis

We first investigated the influence of geography and environmental conditions on the global host population structure of *A. spathulata* across the Western Pacific (N=999). For this purpose, we retrieved environmental predictors from several satellite products and models with a focus on variables known to be primary drivers of coral holobiont community composition (temperature, light and turbidity; Supplementary Material and Methods, Table S5). We modeled spatial relationships among colonies using distance-based Moran’s Eigenvector Maps (dbMEMs), which capture multi-scale spatial relationships using principal coordinate analyses (Borcard & Legendre, 2002). As individual colony-level GPS coordinates were not recorded at NC and CB reefs, randomly jittered coordinates within 300 m around the reef coordinate (recorded using a GPS during sampling; corresponding to the approximate size of the sampling area) were assigned to each colony at these reefs. The neighbor matrix for each colony was computed from spatial coordinates using Haversine distance in R package *geosphere* (Hijmans et al., 2017) and used to compute dbMEMs eigenvectors using R package *adespatial* (Dray et al., 2018). The first four eigenvectors associated with positive eigenvalues were kept in further analyses based on a screeplot of cumulative variance explained (>95%). All predictors were standardized by subtracting the mean and dividing by the standard deviation to ensure comparable units.

We performed a redundancy analysis (RDA; Legendre & Anderson, 1999) using R package *vegan* (Oksanen et al., 2013). The first RDA model was built using the first 10 principal components of the PCA on host SNPs as a dependent matrix, and all environmental variables as predictors. Collinear predictors were removed successively, starting with the variables with the highest variance inflation factor (VIF), recomputing the model VIF and repeating this procedure until all predictors had a VIF < 10. Model significance was tested using an analysis of variance (ANOVA) and individual axes significance was tested using an ANOVA-like permutation test in package *vegan* with 500 permutations per axis. Forward selection was used to select predictors that were statistically significant. We used the framework developed by Capblancq and Forester (2021) to assess the relative influence of geography and environment on host structure. We built a global RDA model using (1) environmental variables previously identified as significant using forward selection and (2) the first four dbMEM eigenvectors as predictors. Separate partial RDA models were built setting either environmental variables or spatial eigenvectors as conditional variables (i.e., ‘partialled out’). Model significance was tested using an ANOVA and models were compared based on inertia, total variance explained, and proportion of constrained variance explained.

We used a similar approach to investigate the influence of geography, environmental factors and host population structure on the differentiation of *A. spathulata* symbiont communities (*n*=878) which are horizontally acquired (van Oppen et al., 2001). Our preliminary analysis showed that ordinations based on *k*-mer-derived D ^s^ distance were influenced by the number of symbionts reads in each sample, therefore all samples were downsampled to 50 M bp and samples below this threshold were discarded (10% of samples). As a small effect on the ordination remained, we included this variable as a conditional variable in all of our models. We performed a principal coordinate analyses (Borcard & Legendre, 2002) on the *k*-mer-derived D ^s^ distance matrix and used the first 10 principal coordinates as a dependent variable in RDA models. As the PCoA did not yield negative eigenvalues indicating the distance matrix was nearly Euclidean, we did not perform negative eigenvalues correction as routinely applied in dbRDA. The first RDA model was built using all environmental variables and significant predictors were selected following the same procedure as above. A second global RDA model was built using (1) environmental variables previously identified as significant using forward selection, (2) the first four dbMEM eigenvectors and (3) the first three host PCs as predictors (explaining almost all of the variation; Figure S8). The marginal effect of each category of predictor was then assessed through separate pRDA models, using the two other categories as conditional variables. All statistical analysis were conducted using *R* v4.0.4 and figures created using package *ggplot2* (Wickham, 2011).

## Results

### Genomic data processing and filtering

We recovered 339,720 single nucleotide polymorphisms (SNPs) from 1,088 individual coral colonies of *Acropora spathulata* using a stringent filtering approach (see Methods). This dataset had a mean locus-level sequencing depth of 13.7 ± 1.2 (mean ± sd). Following removal of related individuals and linked SNPs, Dataset 1 used for population structure analyses comprised 999 individuals and 27,175 SNPs. Dataset 2 used to compute genetic differentiation and diversity metrics consisted of ∼2.2 M loci (SNPs and monomorphic sites). Dataset 3 used for demographic modeling consisted of 37,841–61,902 SNPs per population pair and Dataset 4 used for IbD analyses and *N_e_* estimates consisted of 2,011 SNPs after pruning for physical linkage.

### Coral host population structure

*A. spathulata* colonies across the Western Pacific were separated into four genetically differentiated groups, hereinafter referred to as populations (Figure 1). A PCA revealed geographically distinct populations separated on PC1 and PC2 (1.5% and 0.7% explained variance respectively; Figure 1a), corresponding to the Central to Northern GBR (i.e., from Davies to Hicks reefs), the Southern GBR, the Chesterfields-Bellona atolls (CB) in the Coral Sea and the western New Caledonia’s main island (NC). PC3 (0.5% explained variance, Figure 1a) separated the CB group from the three other groups. Similarly, ADMIXTURE analysis found a concordant optimal number of *K* = 4 ancestral populations (Figure S9, Figure S10). The separation of individuals by their ancestry proportion was consistent with PCA results but revealed small levels of admixture between all populations (Figure 1b). In particular, 4 of 137 and 19 of 137 colonies found in the Southern GBR had a >0.5 and >0.25 assignment to the ancestral population of the Central/Northern GBR respectively.

### Population divergence and genetic diversity

Genetic divergence, estimated using the fixation index, was low between all of the four populations (*F_st_* < 0.030; Table 1) and was the lowest between the two GBR populations (*F_st_* = 0.014). Pairwise *D_xy_* ranged from 0.010 to 0.011 and was the lowest between the NC and CB populations. This corresponds to an average 1% nucleotide difference per bp between populations (Table 1). Nucleotide diversity (π) was the highest in the NC population (π = 0.0106) followed by the Central/Northern GBR (π =0.0103) and the Southern GBR populations (π = 0.0100). The CB population appears to have the lowest nucleotide diversity (π = 0.0097).

**Table 1.** Demographic modeling of *Acropora spathulata* populations divergence across the western Pacific. For each pair of populations (1: left, 2: right), optimized parameters are presented for the model exhibiting the best fit to the observed joint allelic frequency spectrum (divergence with asymmetrical gene flow) along with genetic differentiation statistics (fixation index *F_st_* and absolute divergence *D_xy_*). Nref, *v*_1_, *v*_2_ are the ancestral and current effective population sizes in thousands of individuals. 𝑚 is the proportion of migrants per generation and *M* is the number of migrants per generation. *T* is the estimated divergence time in thousands of years. For each model and parameter, median and 2.5–97.5% confidence intervals based on 100 optimized bootstrap replicates are presented.

| Population pair | Nref | $\nu_1$ | $\nu_2$ | $T$ | $D_{XY}$ |
| --- | --- | --- | --- | --- | --- |
| Central/Northern GBR-Southern GBR | 7.2 [4.1, 10.6] | 63.9 [51.1, 76.3] | 32.5 [25.9, 42.3] | 276 [231, 320] | 0.0102 |
| Central/Northern GBR-NC | 9.2 [5.1, 17.6] | 82.9 [66.9, 118.9] | 25.0 [14.5, 33.6] | 323 [218, 379] | 0.0107 |
| Central/Northern GBR-CB | 4.9 [3.7, 9.9] | 61.9 [51.1, 75.8] | 29.6 [21.6, 36.6] | 354 [292, 373] | 0.0102 |
| Southern GBR-NC | 6.5 [4.6, 10.9] | 82.0 [73.6, 87.9] | 32.8 [28.4, 39.5] | 436 [378, 461] | 0.0107 |
| Southern GBR-CB | 7.1 [4.4, 12.3] | 65.8 [58.1, 74.5] | 34.8 [28.6, 43.9] | 395 [334, 431] | 0.0100 |
| NC-CB | 7.2 [6.6, 9.9] | 86.5 [76.4, 96.7] | 48.8 [35.2, 56.1] | 651 [630, 666] | 0.0104 |
| Population pair | m12 (2→1) | m21 (1→2) | M1→2 ($N_e m_{12}$) | M21 ($N_e m_{21}$) | $F_{st}$ |
| Central/Northern GBR-South GBR | 6.2e-5 [4.6e-5, 8.7e-5] | 2.5e-4 [2.0e-4, 3.0e-4] | 7.9 [6.6, 9.6] | 16.4 [14.7, 18.0] | 0.014 |
| Central/Northern GBR-NC | 2.1e-5 [9.4e-6, 3.5e-5] | 2.0e-4 [1.5e-4, 3.0e-4] | 3.6 [2.3, 4.7] | 9.9 [8.6, 11.0] | 0.025 |
| Central/Northern GBR-CB | 5.4e-5 [3.7e-5, 7.2e-5] | 1.2e-4 [9.2e-5, 1.7e-4] | 6.5 [5.2, 8.0] | 7.1 [6.5, 7.8] | 0.028 |
| Southern GBR-NC | 1.7e-5 [1.3e-5, 2.5e-5] | 1.4e-4 [1.2e-4, 1.5e-4] | 2.8 [2.2, 3.7] | 8.8 [8.2, 9.5] | 0.029 |
| Southern GBR-CB | 5.1e-5 [3.5e-5, 7.1e-5] | 9.3e-5 [5.9e-5, 1.2e-4] | 6.8 [5.1, 8.6] | 6.3 [5.1, 7.3] | 0.026 |
| NC-CB | 6.5e-5 [5.2e-5, 7.3e-5] | 7.0e-5 [5.6e-5, 1.1e-4] | 11.0 [9.8, 12.1] | 6.9 [6.2, 7.8] | 0.026 |

### Dispersal and effective population size

No significant IbD occurred within populations. However, the pattern of IbD was revealed when populations were analyzed jointly, with the regression of individual Rousset’s genetic distance on geographic distance yielding a significant positive slope (β = 0.00064 ± 3.7 × 10^-05^; median ± sd; Figure 2a). The effective population size at the neighborhood size across this geographic range was estimated at *N_e_* = 2,954 ± 1,573 effective individuals (median ± sd; Figure 2b). Combining these estimates, the average dispersal distance between parents and offspring across generations was 109 ± 33 km (mean axial dispersal distance; σ; median ± sd; Figure 2c).

**Figure 2.**
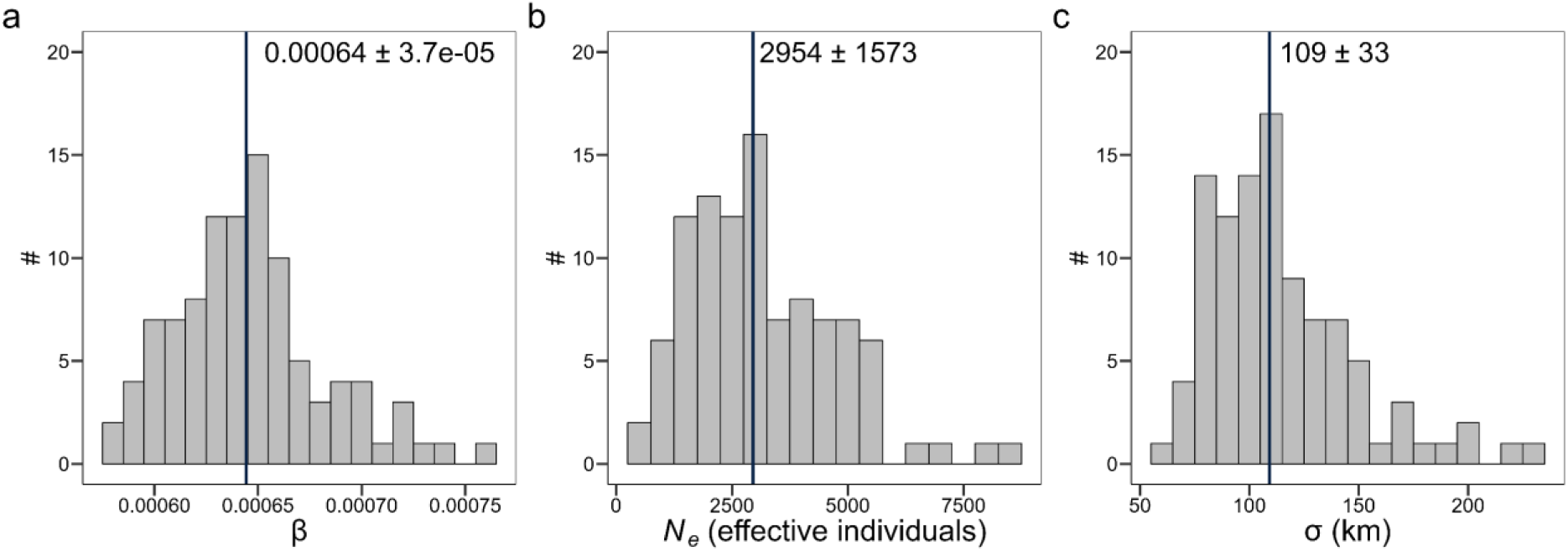
Generational dispersal and effective population size in *Acropora spathulata* from the western Pacific. Isolation by distance at multiple appropriate spatial scales was used to infer the distribution of (a) slope of genetic distance on geographic distance regressions, (b) effective population size at the neighborhood size and (c) mean axial generational dispersal distance. Vertical lines indicate the median of each distribution with median and standard deviation values indicated next to it.

### Demographic modeling of populations divergence and gene flow

For all population pairs, observed joint allelic frequency spectrum (JAFS) were best explained by demographic models consisting of divergence with asymmetrical gene flow than models with no gene flow or ancestral gene flow (Figure S5). This suggests that these populations have diverged across the western Pacific while exchanging larvae over many generations. Demographic models estimated divergence time between 275 to 651 thousand years ago depending on population pairs (Table 1), although these are approximate estimates because germline mutation rates are lacking for this species. Based on these estimates, the two populations from the GBR appear to have diverged most recently (275 [230–319] kya; median [5–95%]). Modelling results suggest that divergence of NC from other populations (323–651 kya) is more ancient than divergence of CB from GBR populations (354–385 kya). However, large overlap in confidence intervals, and possible bias introduced by high migration rates, make it difficult to reconstruct their precise sequential divergence history.

Migration rates and gene flow rates were asymmetrical for most population pairs, and this result was consistent across bootstrap replicates and optimization runs (Table 1). On the GBR, migration rates were on average 5 times higher from the Central/Northern population to the Southern population (2.5 × 10^-04^) than vice versa (6.2 × 10^-05^), resulting in higher southward gene flow (*N_e_*m) in line with predominant oceanographic currents (Figure 1c; Figure S1). Eastward migration rates from both GBR populations to the CB population (9.3 × 10^-05^–1.2 × 10^-04^) were higher than the westward rates (5.4 × 10^-05^–1.2 × 10^-04^). However, differences in long-term effective population sizes (29,000–66,000 effective individuals) resulted in nearly symmetrical gene flow between the GBR and these Coral Sea atolls (6.5–7.1 migrants per generation, Figure 1c). Migration rates between the CB and NC populations were broadly symmetrical (6.5–7.0 × 10^-05^), but the compounding effect of unequal population sizes produced a dominance of eastward gene flow (11.0 [9.8–12.1] migrants per generation; median [5–95%]). Gene flow was also directed primarily eastward, from both GBR populations toward NC (Figure 1c).

### Symbiodiniaceae genetic diversity based on molecular markers

We mapped non-coral reads from our 1,088 hologenomes to Symbiodiniaceae reference genomes and recovered a median of 708,000 Symbiodiniaceae reads per sample (range: 63,540 to 24,932,887). All colonies harbored symbionts from the *Cladocopium* genus with predominant ITS2 sequence variants (recovered from hologenome reads) classified as C50, C3 and *Cladocopium* sp. (Figure S11). Hologenome sequencing results were consistent with amplicon sequencing results in a subset of 36 samples, although the latter showed a greater taxonomic resolution, separating profiles dominated by C3k, C3k/C3bo, C50b, and C50c sequences (Figure S6). Read mapping against *psbA*^ncr^ sequences revealed classifications of *C. vulgare* (C1) or *C. sodalum* (C3; Figure S11). The mapping of reads to *psbA*^ncr^ C1 in samples associated with C50 ITS2 sequences was presumably due to the lack of reference *psbA*^ncr^ sequences of *Cladocopium* C50, limiting the use of this method for finer taxonomic classification. However, our results suggest that the C3 symbionts identified are more closely related to *C. sodalum* than other described C3 species (>80% of mapped sequences; Butler et al., 2023).

### Symbiodiniaceae community divergence based on k-mer profiles

Alignment-free clustering of symbiont reads revealed some extent of co-clustering between each of the four main symbiont taxa and the corresponding host populations (Figure 3). The host population in the Central/Northern GBR was associated predominantly with *Cladocopium* taxa from C3k and C50c subclades, with a cross-shore partitioning of these communities. Offshore reefs were associated with C3k, while inshore reefs were associated with C50c (Figure 4a). At intermediate reefs (e.g., Moore, North Direction, Mackay), some colonies harbored the C50c taxon while others harbored C3k. The host population in the Southern GBR associated with a symbiont taxon of the C50b subclade. A few colonies associated with C3k at Lady Musgrave Island. Corals in Western NC were predominantly associated with a taxon from the C3k/C3bo subclade. The exception was some colonies in most southern reefs which associated with a distinct symbiont taxon from C50b subclade. Based on ITS2 profiles, *k*-mer profiles clustering and geographic separation, this symbiont taxon is likely genetically differentiated from the C50b taxa found in the Southern GBR (Figure S6 and S7). Coral hosts in the CB atolls were also associated with a C3k/C3bo taxon that was indistinguishable from NC C3k/C3bo despite being separated by >400 kilometers.

**Figure 3.**
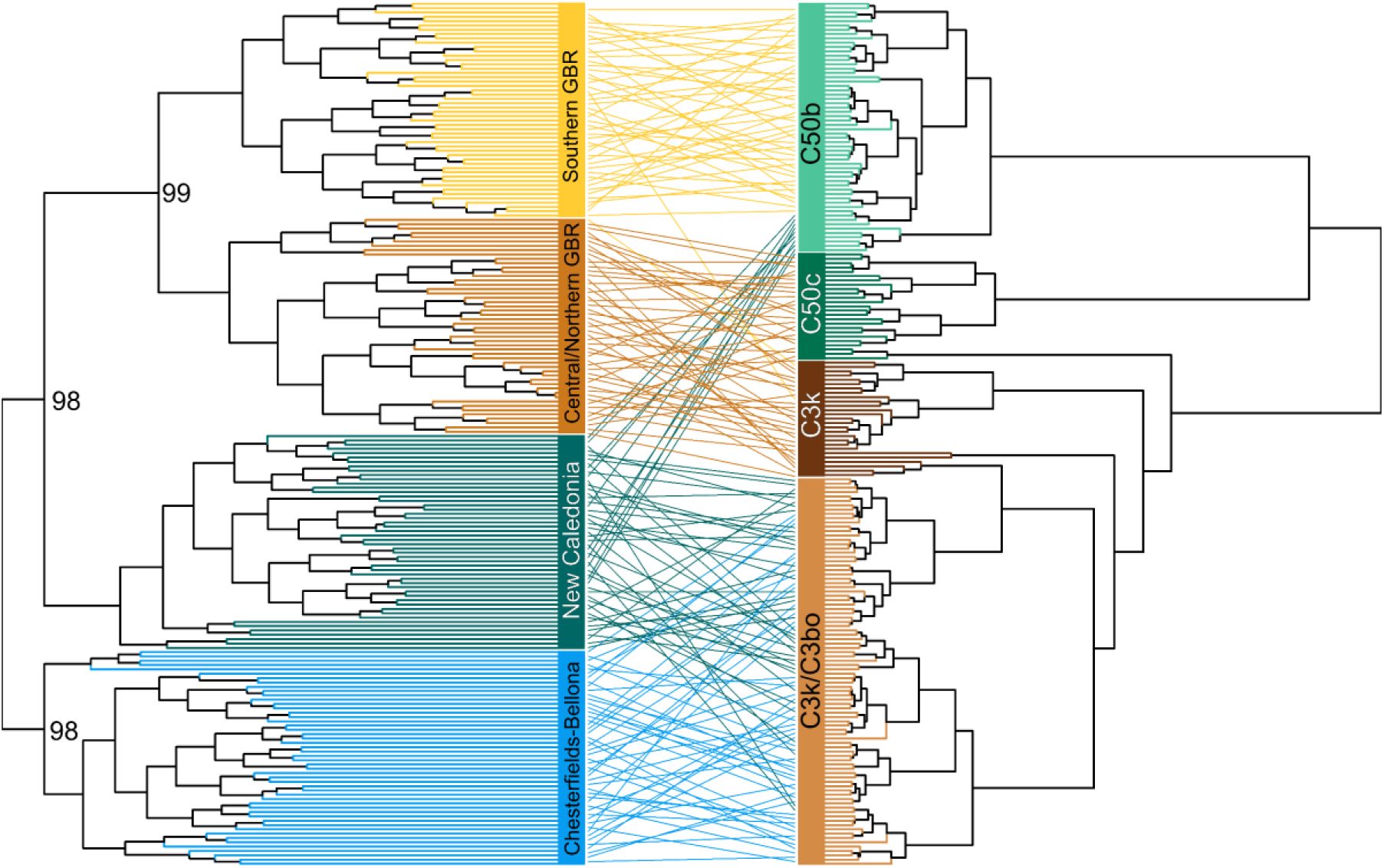
Co-clustering of *Acropora spathulata* coral hosts and associated Symbiodiniaceae taxa across the western Pacific. The host dendrogram (left) is a maximum likelihood tree derived from 27,175 SNPs using RAxML. The node values represent bootstrap support (>90) from 100 bootstrap replicates and branches are coloured based on ADMIXTURE ancestral populations. The symbiont dendrogram (right) is obtained through hierarchical clustering of the D ^s^ distance matrix based on symbiont *k*-mer profiles and branches are colored according to majority ITS2 sequences for each sample.

**Figure 4.**
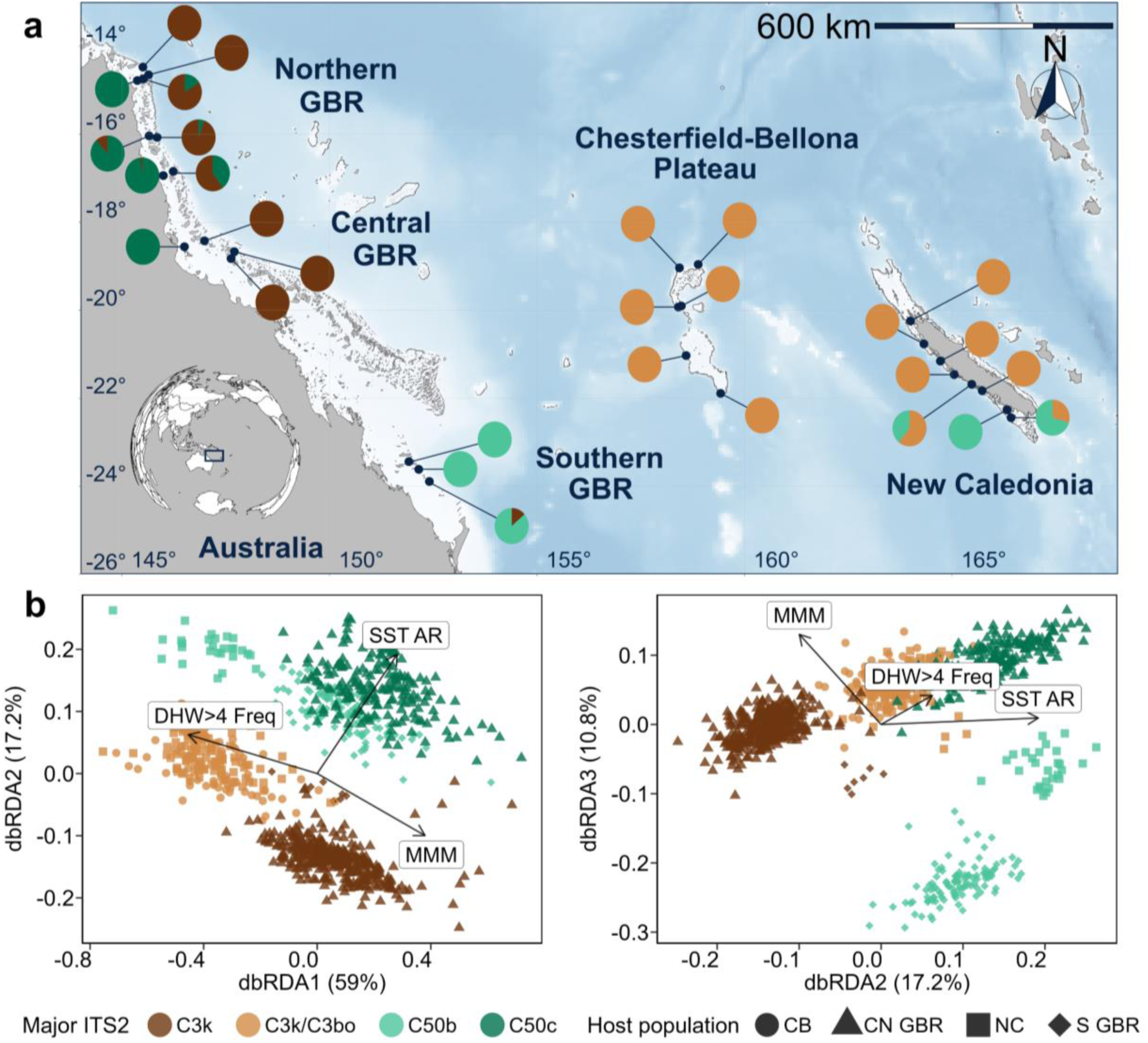
Variation in the symbiont (Symbiodiniaceae) communities of *Acropora spathulata* across geographic and environmental gradients in the western Pacific. (a) Map showing the proportion of colonies harboring each of the symbiont clusters identified using ordination of D ^s^ distance matrix at each sampling site. Symbiont genetic clusters are coloured and labelled based on their major ITS2 type. (b) Distance-based redundancy Analysis (db-RDA) ordination of D ^s^ *k*-mer profiles. Each individual is coloured by its major ITS2 sequence and the shape corresponds to each of the host populations (CN GBR = Central/Northern GBR, S GBR = Southern GBR, NC = New Caledonia, CB = Chesterfields-Bellona). Arrows represent significant environmental variables in the model. The first three axes explaining 87% of the constrained variance are shown.

### Drivers of host and symbiont genetic structure

The RDA indicated that environmental variables significantly predicted coral host genetic structure (ANOVA; *p*-value < 0.01), explaining 64% of the total variance. Maximum monthly mean (MMM) sea temperature, the frequency of mild marine heatwaves (DHW > 4), and distance to coast were the strongest reef-level environmental predictors of host population structure (cumulative adjusted *R^2^*= 0.60, Table 2). A global RDA including geographic predictors (dbMEMs) on top of significant environmental variables explained 70% of the total variance in host genetic structure. Due to the spatial sampling design, several partial-RDAs indicated that environmental and geographic effects were largely confounded (86% of explained variance).

**Table 2.**
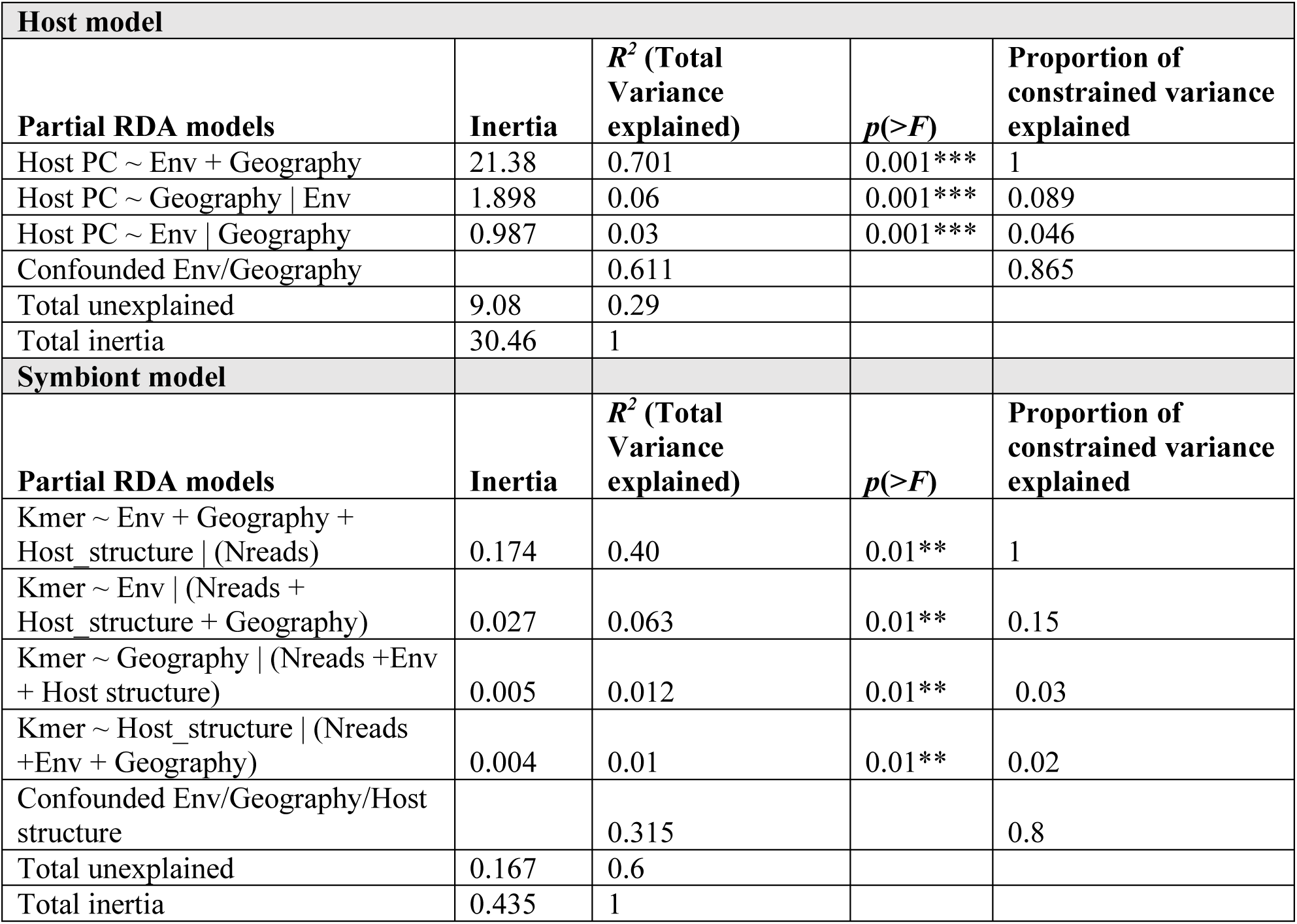
The influence of geography and environmental conditions on host neutral genetic structure of *Acropora spathulata* and the influence of geography, environmental conditions and host structure on associated Symbiodiniaceae communities. The relative influence of each category of predictors was decomposed using partial redundancy analysis (pRDA). Geography was modeled using distance-based Moran’s Eigenvector maps. Environmental variables included factors indicative of temperature, irradiance, and turbidity.

Environmental predictors also significantly influenced the community composition of *Cladocopium* symbionts (ANOVA; *p*-value < 0.01), explaining 37% of the total variance. The maximum monthly Mean (MMM), frequency of mild marine heatwaves (DHW > 4) and annual range in sea surface temperature (AR) were retained as significant predictors in the model (*p*-value = 0.001). The AR separated C3 populations, found in more thermally variable environments at inshore and high latitude reefs, from C50 populations found in offshore and low-latitude reefs (Figure 4b). Within the C50 subclade, distinct C50b and C50c populations were separated primarily by the MMM reflecting both temperature variation across latitudinal gradient and cross-shore gradients. Within the C3 subclade, the C3k populations were positively associated with the MMM, while C3k/C3bo populations were found in reefs where MMM was lower on average, but where mild marine heatwaves have been more frequent. Several pRDAs indicated that the environment was the strongest driver of symbiont community composition (6.3% of total explained variance). Geography (1.2%) and host population structure (1%) were less influential, although the effect of these three components was also largely confounded (80% of explained variance, Table 2).

## Discussion

Out results revealed four genetically distinct *A. spathulata* populations in the studied western Pacific region, which have diverged due to geographic isolation but maintain gene flow through major oceanographic currents. These populations show little genetic structure over distances greater than 500 km, reflecting a generational dispersal distance estimated at ∼100 km. In contrast, coral hosts are associated with distinct algal symbionts of five Symbiodiniaceae taxa across shorter latitudinal and cross-shelf gradients. This study presents one of the most extensive datasets for analyzing spatial genetic variation in both corals and their algal symbionts using whole-genome sequences, combining broad geographic coverage and dense local sampling. These results suggest that large local populations and dispersal support resilience of a broadcast-spawning coral, whereas habitat-specific symbionts and long-distance gene flow could enhance its adaptive potential in changing environments.

### Regional spatial genetic variation differs among holobiont partners

The spatial scale of genetic variation differed markedly between *A. spathulata* and its associated *Symbiodiniaceae*. This coral species comprised four genetically differentiated populations distributed in distinct regions separated by ∼600–2,200 km. However, each population was largely genetically homogenous based on PCA, ADMIXTURE and isolation-by-distance analyses, despite spanning up to >300km for NC and CB, and up to >500km for the Central/Northern GBR. Conversely, Symbiodiniaceae community genetic variation occurred over spatial scales as small as 30km, and colonies from the same reef sometimes hosted distinct symbiont taxa. These patterns reflect the predominant effect of distinct evolutionary forces acting on holobiont partners.

Patterns of *A. spathulata* genetic variation suggest extensive gene flow promotes genetic homogenization of populations, counteracting the effects of drift and local adaptation (Slatkin, 1987). This was supported by our estimated σ exceeding >100 km. Across the distant western Pacific populations, reduced gene flow and increasing environmental contrasts (e.g., Δ∼3°C in MMM; Figure S1) drive greater genetic differentiation (*D_xy_* = 0.0102–0.0107). However, the relative importance of genetic drift and local adaptation in this differentiation could not be disentangled due to spatial confounding between environmental and geographic predictors (Table 2).

Patterns of Symbiodiniaceae genetic variation suggest that environmental filtering is the predominant driver of *A. spathulata* algal symbioses. Variation in symbiont communities was likely not strongly influenced by dispersal, as it did not follow an IbD pattern. For instance, colonies from Martin and No Name reefs (30km apart) hosted different taxa, while those from No Name and Chicken reefs hosted the same taxa (500km apart; Figure 4). In addition, the environment had a larger effect on symbioses than host genetic differentiation (although largely confounded; pRDA; Table 2), as expected from species that acquire symbionts from their environment after settlement (Davies et al., 2020; van Oppen et al., 2001). Variation in symbioses was particularly pronounced across shelf and latitudinal gradients, similar to other *Acropora* species on the GBR (Cooke et al., 2020; LaJeunesse et al., 2004; Matias et al., 2022). Notably, *A. spathulata* and *A. tersa* (formerly *A. hyacinthus*) hosted strikingly similar Symbiodiniaceae communities, with identical ITS2 types found across comparable environmental gradients (Naugle et al., 2024). RDA suggest that temperature was the primary driver, although other important factors such as light and turbidity (Cooper et al., 2011) were likely excluded due to multicollinearity. It remains also unclear whether *A. spathulata* associates with specific Symbiodiniaceae because these taxa are the most abundant at these locations, because they form the best performing symbioses (Bhattacharya et al., 2024), or both—since their relative abundance in the environment is unknown.

The spatial distribution of Symbiodiniaceae suggests some taxa may be environmental specialists while others may be environmental generalists. Similar to Epstein et al. (2019), we found *A. spathulata* associated with a C3k taxon at cooler offshore Central/Northern GBR reefs, and we also revealed a distinct C50c taxon predominant at warmer turbid inshore Central/Northern reefs. Conversely, C3k/C3bo appears to have a wide distribution as it was indistinguishable based on genome-sequence composition between NC and CB. Taxa from the C50b subclade, found across a 13° longitudinal gradient, might also be widespread in this region but best performing at high-latitudes.

Importantly, these findings show that phylogenetic relatedness does not necessarily imply similar ecological niches, as the C50c and C50b populations were found in warmer and cooler environments respectively.

### Broadcast spawners maintain long-distance asymmetrical gene flow across the Western Pacific

Demographic models detected substantial historical gene flow between Western Pacific populations separated by thousands of kilometers, that likely started to diverge between 0.275 to 0.651 MYA. While this potential connectivity had been suggested by biophysical models of coral dispersal (Wood et al., 2014), it remained to be empirically tested due to their potential biases (Saint-Amand et al., 2023). Although some uncertainty remained in the absolute values of gene flow, primary directions of migration rates were consistent across bootstrap replicates for all population pairs and these models had stronger support than divergence without gene flow.

On the GBR, gene flow was primarily southward, consistent with the predominant EAC direction and other studies on coral broadcast spawners (Matz et al., 2018; Meziere et al., 2025; Riginos et al., 2019). We also observed admixture between the Central/Northern and Southern GBR populations indicating recent southward migration between these regions (Figure 1). Conversely, gene flow was largely symmetrical between the GBR and Coral Sea atolls, reflecting the region’s complex oceanography, which includes westward currents such as the North Caledonia Jet (NCJ) and the South Caledonia Jet (SCJ; Cravatte et al., 2015), as well as the eastward Sub Tropical Counter Current (STCC; Marchesiello et al., 2010). Demographic models inferred dominant eastward gene flow from the GBR to NC, and from CB to NC, which is unexpected given the prevailing westward NCJ and SCJ in the region (Cravatte et al., 2015). These patterns of realized connectivity may have been influenced by surface circulation associated with the STCC, while the strong NCJ and SCJ reach their maximum intensity in deeper layers (Marchesiello et al., 2010). Observations of plastic debris and Lagrangian dispersal modeling notably suggest that mesoscale currents can generate eastward drift in the region despite the dominance of westward zonal jets (Maes & Blanke, 2015). Because gene flow inferred from demographic models reflects an average across thousands of generations, these estimates could also have been influenced by temporal changes in ocean circulation. Together, these results provide novel insights into a complex realized connectivity across this region and highlight the value of such analyses for informing management in contexts of variable ocean circulation. Beyond uncertainties in the current directionality of gene flow, the models strongly support low but persistent migration between these western Pacific populations. As such, they should be treated as distinct management units (Hohenlohe et al., 2021) within a broader regional network that also recognizes the importance of their connectivity for evolutionary adaptive potential.

### Large breeding population and high dispersal may support demographic resilience

The effective population size of *A. spathulata* at the neighorhood size was relatively high (*N_e_*=2,950) despite drastic coral mortality over last decades (Eddy et al., 2021; Souter et al., 2021). This estimate is particularly important for its conservation, as emphasized by the recent Kunming-Montreal Global Biodiversity Framework (Waples, 2024). Although there is no exact rule of how large *N_e_* should be, our estimate suggests that local breeding populations are sufficiently large to prevent dramatic loss of genetic diversity due to genetic drift (Waples, 2024). In contrast, brooding species with limited dispersal and local recruitment tend to have lower *N_e_* and higher local extinction risk (Hernández-Agreda et al., 2024; Meziere et al., 2025; Prata et al., 2024).

High *N_e_* likely reflects substantial census sizes of this common species densely distributed on reef flats and slopes, combined with a large generation dispersal estimated at ∼100km per generation. This estimate contains some level of uncertainty because IbD theory assumes symmetrical dispersal across a continuous habitat of constant population density, which does not match the typical patchiness of coral reefs. Yet, this value is consistent with a pelagic larval duration estimated from lipid reserves at ∼100 days which would allow larvae to be carried across large distances (Graham et al., 2013). Estimates from previous studies using similar methods vary from a few meters in brooding species (Gorospe & Karl, 2013; Hernández-Agreda et al., 2024; Prata et al., 2024) to 1–52 kilometers in broadcast spawning species (Japaud et al., 2019; Meziere et al., 2025). Our estimate also falls at the upper end of broadcast-spawners connectivity predicted by biophysical models (63% of settlement within 100km; Wood et al., 2014) and is comparable to coral trout dispersal on the GBR (Williamson et al., 2016). These findings suggest that *A. spathulata* exhibits one of the highest coral dispersal distances recorded to date. Importantly, we only estimated a global isotropic σ because IbD was not significant within populations but this estimate could vary among regions. Even if σ is similar across regions, the demographic consequences could differ because the GBR is a fragmented mosaic of numerous discrete reefs while NC and CB contain longer continuous barrier reefs (Andréfouët et al., 2009). Individual reefs are expected to be demographically isolated when separated by more than 2σ (Pinsky et al., 2012) and thus connectivity could differ between these reef systems of contrasting geomorphology.

Large *N_e_* and high dispersal may explain the impressive recovery of *Acropora* communities on many GBR reefs after the 2016–2017 bleaching-related mortality events (Emslie et al., 2024), and will be critical for demographic recovery under increasingly frequent disturbances (Hock et al., 2017). For example, marine heat waves and freshwater floods caused 90–100% mortality of *A. spathulata* colonies at several inshore Northern GBR sites sampled in this study in 2024, while colonies at offshore reefs were relatively unaffected. Larval dispersal from these populations could thus in theory supply larvae to degraded reefs. However, a finer assessment of realized dispersal is needed to evaluate the inbound and outbound connectivity of individual reefs, as it depends on meso-scale currents and habitat availability. Finally, given the ongoing impacts to these populations from global warming, monitoring possible changes in *N_e_* will be important to track changes in their vulnerability (Howells et al., 2022).

### Habitat-specific symbioses and long-range dispersal affect the evolutionary adaptive potential of A. spathulata

*A. spathulata* formed distinct symbioses with Symbiodiniaceae across distinct thermal conditions which raises the question of whether shifts to more heat-tolerant symbioses could promote the adaptation of *A. spathulata* to rapid climate warming. *Acropora* corals symbioses with heat tolerant symbionts—notably from the *Durusdinium* genus—have been observed to increase in prevalence during marine heatwaves but this response appears to be species-specific and has not been yet reported for *A. spathulata* (Quigley et al., 2022). The absence of *Durusdinium* in colonies sampled outside major marine heatwaves in Epstein et al. (2019) and in our study suggests this association is uncommon in *A. spathulata* under normal conditions. However, we observed variable symbioses among colonies at mid-shelf reefs on the GBR, suggesting that both the C50c and C3k taxa are found in these environments. Under future climate warming, symbioses with C50c typically found in warmer inshore environments may become more prevalent on mid-shelf reefs, especially given that fast growing *Acropora* corals can rapidly shift to better adapted symbionts within 1–2 years (Turnham et al., 2025). Whether this ability varies across regions remains an open question, especially given that the same C3k/C3bo taxon was found in colonies both from NC and CB reefs despite drastically different environments. The absence of other associations may indicate lower potential for adaptation through shifts in symbioses, or on the contrary indicate greater physiological plasticity and resilience to changing conditions of this symbiont taxon. Further sampling around bleaching events is thus needed to assess whether this species can shift symbioses quickly enough to reduce coral mortality over future marine heatwaves.

Another factor that may underlie the evolutionary adaptation of *A. spathulata is the* long-distance gene flow between western Pacific populations, as it could increase genetic variation or introduce heat-tolerant alleles from warmer to cooler regions (Matz et al., 2020). On the GBR, dominant southward gene flow may allow this process, as observed in other coral taxa (Matz et al., 2018; Meziere et al., 2025; Riginos et al., 2019), particularly given that admixture suggest recent southward migration. Among other populations, the effect of gene flow on adaptive genetic variation may be harder to predict due to more-complex patterns. For example, the southern GBR showed substantial northeastward gene flow toward CB populations typically found in warmer environments. Nevertheless, occasional long-distance gene flow will benefit populations’ adaptive potential by introducing novel genetic variation without swamping adaptive local variants (Fitzpatrick et al., 2020). We only inferred broad-scale patterns of gene flow among the four regions. Sampling a continuum of reefs in these regions and modeling gene flow at a finer resolution among reef groups (e.g., Matz et al., 2018; Meziere et al., 2025), along with construction of high-resolution biophysical models (e.g., Mason et al., 2025), would help building finer networks of reef connectivity (Mumby et al., 2021).

Within regions, high level of gene flow will constrain evolution by preventing strong local adaptation, for instance along cross-shore gradients. However, it will also facilitate the spread of standing genetic variation, as evidenced by limited differences in genetic diversity between reefs and regions. This gene flow will reduce the risk of inbreeding depression and enable the rapid spread of novel genetic variants across the population range (Slatkin, 1987). Future work investigating the distribution of adaptive genetic variation in this species will help clarify this ambivalent effect.

## Conclusion

The genetic structure of the Western Pacific coral *A. spathulata* reflects opposing forces of geographic isolation and long-distance dispersal driven by major regional ocean currents. In parallel, variation in associated *Cladocopium* taxa along latitudinal and cross-shore environmental gradients suggests environmental filtering of symbioses. Our results enable the definition of management units across the Western Pacific, and emphasize the importance of considering coral reefs connectivity at the regional scale beyond national borders—as maintaining gene flow between distant populations could underlie their evolutionary adaptation (Fitzpatrick et al., 2020). The distinct spatial scales at which evolutionary processes affect coral hosts and their symbionts are also essential to consider for many proposed intervention approaches such as assisted gene flow, selective breeding or symbiont manipulations (van Oppen et al., 2017). Our findings indicate substantial effective population size and exceptional dispersal in *A. spathulata* that may support demographic rescue following disturbances, provided some locations remain unaffected. However, this study only offers a snapshot of the demographic status of *A. spathulata* populations. In light of the recent severe coral mortality events (Cantin et al., 2024), future reassessment of their vulnerability will be essential for effective conservation.

## Resource availability

### Lead Contact

Requests for further information and resources should be directed to and will be fulfilled by the lead contact, Hugo Denis.

### Data and code availability

Raw sequence data and unfiltered VCF file from this study will be made available on NCBI upon publication (BioProject XXX and BioProject XXX) and associated sample metadata can be accessed on GEOME (https://geome-db.org/record/ark:/21547/GPk2 and https://geome-db.org/record/ark:/21547/GLz2). Scripts used for WGS data processing and analyses are available on GitHub (https://github.com/hde08/Aspat_CoralSea_Popgen) and will be archived on Zenodo upon publication (XXX). Any additional information required to reanalyze the data reported in this paper is available from the lead contact upon request.

## Acknowledgements

We thank the Traditional Owners of the Great Barrier Reef and New Caledonia reefs for their free prior and informed consent to undertake coral sampling in their Sea Country. We thank the government of New Caledonia for granting permission to conduct coral sampling within the Natural Park of the Coral Sea, as well as the South Province and North Province. We thank Samantha Goyen, Riverside Marine Cruise crew members and numerous AIMS staff and volunteers for their support while collecting on the GBR. We thank the French National Navy, Mahé Dumas, Bertrand Bourgeois, William Roman, Jordi Giraud and Martin Nguyen for their help while collecting in Chesterfield-Bellona atolls and New Caledonia Grande Terre. We thank Samantha Howitt for assisting in the DNA extractions at the University of Queensland and Clarisse Majorel for assisting in the DNA extractions in New Caledonia. We thank Molly Przeworski for providing an upgraded reference genome assembly. We thank Jérôme Lefèvre and Christophe Charbuillet for their support with HPC systems. This research was undertaken with the assistance of resources from the National Computational Infrastructure (NCI Australia), an NCRIS-enabled capability supported by the Australian Government, through Project d85 awarded to C.X.C. Finally, we are thankful to Ryan Gutenkunst for his advice on demographic modeling analyses.

## Authors contributions

V.B.L., E.J.H., L.K.B., C.R. and H.D. initiated the project and planned the sampling campaign. Fieldwork was done by H.D., L.K.B., V.M. and E.H. on the GBR and H.D., V.B.L., M.B. and G.L. in New Caledonia. DNA extractions were performed by I.B. and K.E.P. for GBR samples, and by H.D for New Caledonia samples. Whole genome sequence data was curated and analyzed by H.D. following a pipeline designed by I.P. Host population structure and phylogenetic analyses were performed by H.D. supported by I.P, K.E.P., C.R., E.J.H., G.L. and V.B.L. Demographic modeling analyses were performed by H.D. supported by K.E.P. IbD analyses were performed by K.E.P. Analyses of symbiont communities were performed by H.D. supported by H.I and C.X.C. H.D. wrote the first draft version of the manuscript which was critically revised by all co-authors. All authors read and approved the final manuscript.

