## Supplemental materials for "Long-distance gene flow and contrasting population structures of reef-building corals and their algal symbionts inform adaptive potential across the Western Pacific"

### Supplementary material

### Methods and Results

#### *Testing for batch effects*

For logistical and legal reasons, samples collected in Australia and New Caledonia were processed for DNA extraction, library preparation and sequencing at different facilities. While we used similar DNA extraction kits, library preparation kits and sequencing platform to minimize batch effects in our dataset, a subset of 23 GBR samples were extracted and sequenced separately at both facilities to validate the absence of strong batch effects. Following the different data processing steps described in the main manuscript, 17 pairs of samples passed filtering steps and were used to assess the presence of batch effects using 'hard called' variants. First, each pair of replicates were correctly grouped together based on relatedness, irrespective of the batch in which they were sequenced (Supplementary Figure S3a). Second a PCA conducted on SNPs (MAF > 0.01, missingness < 20%) correctly separated GBR samples in the two previously described genomic clusters, rather than by batch (Supplementary Figure S3b). Third, we computed individual autosomal heterozygosity for each batch using 0% missing data. This was done because different factors such as different levels of DNA degradation leading to base transition can alter heterozygosity estimates between batches (Lou & Therkildsen, 2022). A Welch two-sample t-test found no significant differences in heterozygosity between batches. Overall, these results confirm the absence of strong batch effects in our dataset. While slight differences between samples extracted and sequenced separately are to be expected, the genetic patterns described in this study are unlikely to be an artifact of batch effects.

#### *Choice of reference genome*

Cleaned reads were initially aligned to *Acropora millepora* v3 reference genome (unpublished upgrade from GCA_013753865.1_Amil_v2.1). This reference was chosen as *A. millepora* is a very close taxa to *A.spathulata*, both phylogenetically and morphologically (Souter et al., 2010) and had a curated chromosome-scale reference genome at the time analyses were performed unlike *A. spathulata*. Since initial analyses were performed, two *A. spathulata* reference genomes have become available on NCBI Genbank (one chromosome scale; GCA_964019555.1 and one contig-scale; GCA_031770025.1). However, alignment of both reference genomes using D-GENIES (Cabanettes & Klopp, 2018) revealed a moderate macrosynteny between the two *A. spathulata* reference genomes and limited microsynteny with less than 5% of the reference genomes showing > 75% identity. For comparison, these levels of identity or on the same order of magnitude than between two different species: *A. tenuis* with *A.millepora* (Cooke et al., 2020) which suggests that better curation of these references might be needed. In addition, preliminary tests mapping reads from our samples to the contig-scale *A. spathulala* reference (GCA_031770025.1) and *A. millepora v3* showed only a < 0.3% difference in mapping rates between them. The better quality, curation and assembly scale of *A. millepora v3* reference genome thus made it a preferable choice for our analyses.

*Host population genetics using genotype likelihoods*

To get an initial impression of the population structure across our study system we analyzed coral samples genomic data with a genotype likelihood framework in ANGSD v0.940 (Korneliussen et al., 2014). The dataset comprised the 1,132 putative *A. spathulata* samples and validated *A.millepora* (n=13) samples used as outgroups, processed using the same pipeline. Polymorphic sites were filtered to retain only those with mapping quality > 30, base quality > 30, coverage ≥ 3 paired-end reads in ≥95% of individuals, and sites mapping to 14 assembled chromosomes. Genotype likelihoods were inferred using Samtools model to estimate major and minor alleles frequencies assuming biallelic sites (-doMaf = 2) and considered only polymorphic sites with a likelihood ratio test p-value < 0.000001.

*A.spathulata* is a formally recognized species on the GBR. A first subset of the dataset including 864 GBR individuals was used to pinpoint potential taxon mis-identification. To that end, polymorphic sites with minor allele frequency (MAF) > 0.05 were used to compute a covariance matrix and perform a principal component analysis (PCA) in PCAngsd v1.21 (Meisner & Albrechtsen, 2018). Bayesian hierarchical clustering admixture analyses was also performed in PCAngsd with the 'admix' option assuming 2–5 ancestral populations. Both analyses revealed two major genomic clusters geographically separated between the central/north and south GBR. In both analyses, 4 samples were also detected as being *A.millepora* colonies that were mis-identified in the field and were discarded in further analyses (Supplementary Figure S15).

*A.spathulata* has been less often identified and studied in New Caledonia despite being reported to occur in the Western Lagoon of the main island 'Grande Terre' (Fenner & Muir, 2008). Prior work targeting *A.millepora* has also revealed several genetic clusters that could actually consist of the two species or other related taxa (Selmoni et al., 2021). In the field, colonies most closely matching *A.spathulata* morphology from the Great Barrier Reef were targeted. We further assessed how these colonies related to the GBR *A.spathulata* and *A.millepora* by repeating the same PCA and Ancestry analyses using a dataset of 1,041 colonies from the Central/North and South GBR, New Caledonia as well as *A.millepora* from the central GBR (Supplementary Figure S16). These analyses suggested that colonies collected in New Caledonia were indeed most closely related to the GBR *A.spathulata* than the GBR *A.millepora*. Colonies sampled in the western coast of New Caledonia (NC) and the Chesterfields-Bellona plateau (CB) formed two genetically distinct groups. In addition, a small subset of 23 samples from the southern west coast of New Caledonia clustered separately from the rest, presumably originating from an undersampled genetically distinct group that may be closer to the GBR *A.millepora*.

*Validation of genetic differentiation from A.millepora using hard called variants*

To confirm genetic differentiation of previously described groups, an even subset of 11 individuals for each of the 6 groups (*GBR A.millepora*, Central/North GBR *A.spathulata*, South GBR *A.spathulata*, NC Cluster 1, NC Cluster 2 and CB) was used for SNPs and variant calling using the same GATK pipeline and filtering steps as described in the main manuscript. The resulting vcf file was used to compute Pairwise $F_{st}$ between each group using the Weir and Cockerham method (Weir & Cockerham, 1984) in *vcftools*, and used to perform a PCA with the *glPca* function from R package *adegenet* v2.1.10 (Jombart, 2008). These analyses confirmed that all sampled taxa in this study were most closely related to the GBR *A.spathulata* (Supplementary Figure S17). The PCA analysis also confirmed the presence of two genetically distinct groups in the western coast of New Caledonia. Due to its low sample size, the under sampled Cluster 2 was discarded from further analyses to focus on the predominant Cluster 1.

*Clones and related individuals' identification*

Clonemates and related individuals were identified in the dataset using two methods. First, identity-by-state (IBS) matrices were generated in *ANGSD* using the *-doIBS* function from the set of filtered polymorphic sites. Second, relatedness matrices were generated from hard called variants (20% missingness) in *vcftools* using the method of Yang et al. (2010). In both cases, matrices were computed and clonemates identified separately for each of the four genetic groups (NC, CB, South GBR and Central/North GBR). Colonies genetic similarity was compared to the overall background similarity as well as the similarity of technical replicates to infer pairs or groups of clonemates. Both methods were in agreement for all pairs or groups of clones. Using a conservative relatedness threshold of 0.2, there were 11, 17, 3 and 6 groups of clones or related individuals for the Central/North GBR, the South GBR, NC and CB respectively (Supplementary Figures S8 and S9). The number of putative clones and highly related individuals was particularly high in the Southern GBR but we could not confirm if this was due to higher inbreeding rates at these locations or an artifact of the sampling scheme, as colonies were sampled randomly by different groups of divers between locations. All related individuals identified using a conservative threshold were filtered prior to final population structure, demographic and isolation by distance analyses to retain one individual with the lowest missingness per group.

*Environmental data acquisition*

To assess environmental conditions driving symbiont community composition across coral host populations and our study range, we retrieved environmental data from several satellite products and model reanalysis (Supplementary Table S5). These datasets were preferred to more fine-scale resolution models available in each country for consistency reasons. We focused on variables that are known to affect Symbiodiniaceae communities structuring and can be relatively well estimated in both reef systems. Temperature metrics were computed from the Coral Watch v3.1 sea surface temperature (SST) satellite product (NOAA; 1985–present; 5km). Light intensity metrics (Cloud fraction and UV radiation) were computed from the ERA5 reanalysis (Copernicus marine service; 2002–present; 0.25°). Water turbidity ($Kd_{490nm}$) and chlorophyll content metrics were computed from ESA Globcolour satellite product (Fanton d'Andon et al., 2009; 2002–present; 1/24°). Finally, we included spatial descriptors in the analysis such as Latitude and the shortest haversine distance to the shore.

### Supplementary Tables

**Table S1.** Summary of sampling effort across coral reef sites in the western Pacific. Maximum Monthly Mean (MMM) of each reef was computed from the Coral Reef Watch of the National Oceanic and Atmospheric Administration (NOAA Coralwatch v3.1 satellite product). Colonies depth was standardized to the tide at the time of collection and are reported relatively to the Lowest Astronomical Tide (LAT). Negative values indicate shallow colonies above the LAT.

| Country | locationID | location Code | Sampling Date | Lon | Lat | MMM (°C) | Depth mean [range] | N |
|---|---|---|---|---|---|---|---|---|
| Australia | Hicks | HICK | 28/02/2022 | 145.502542 | -14.475094 | 28.6 | 2.2 [0.7,4.7] | 61 |
| Australia | North Direction | NDIR | 01/03/2022 | 145.5133 | -14.7434 | 28.6 | -0.2 [-0.7,0.1] | 52 |
| Australia | Martin | MART | 02/03/2022 | 145.3681 | -14.7801 | 28.6 | 0.6 [-0.2,1.2] | 60 |
| Australia | No Name | NONA | 03/03/2022 | 145.6383 | -14.6508 | 28.6 | 1.1 [-0.2,3.6] | 53 |
| Australia | Mackay | MACK | 04/03/2022 | 145.6514 | -16.0384 | 28.8 | -0.2 [-0.9,1.1] | 60 |
| Australia | St Crispin | STCR | 05/03/2022 | 145.8449 | -16.0724 | 28.7 | 0.1 [-0.9,3.5] | 60 |
| Australia | Fitzroy Island | FITI | 06/03/2022 | 145.9948 | -16.9406 | 28.8 | 0.3 [-0.6,2] | 60 |
| Australia | Moore | ONMO | 07/03/2022 | 146.2316 | -16.8457 | 28.6 | 0.2 [-0.3,1.1] | 44 |
| Australia | Pelorus | PAPE | 08/03/2022 | 146.5023 | -18.5587 | 28.7 | 2.1 [0.1,3.3] | 45 |
| Australia | Kelso | KELS | 09/03/2022 | 146.9849 | -18.424 | 28.4 | 0.9 [0.5,5.3] | 44 |
| Australia | Chicken | OCCH | 10/03/2022 | 147.7064 | -18.6706 | 28.4 | 0.9 [-0.4,4.6] | 60 |
| Australia | Davies | OCDA | 12/03/2022 | 147.6268 | -18.827 | 28.4 | 1.2 [0,3.9] | 53 |
| Australia | Lady Musgrave | CBLM | 24/03/2022 | 152.4064 | -23.8929 | 27.1 | 0.9 [-1.1,4.2] | 56 |
| Australia | Fitzroy Reef | FITR | 25/03/2022 | 152.15 | -23.617 | 27.1 | -0.6 [-0.8,0.7] | 61 |
| Australia | Heron | CBHE | 26/03/2022 | 151.912971 | -23.441601 | 27.1 | -0.7 [-1.3,1] | 60 |
| New Caledonia | Bellona Sud | CB1 | 23/06/2023 | 159.425853 | -21.8905 | 26.6 | -0.2 [-0.5,0] | 4 |
| New Caledonia | Bellona Ouest | CB2 | 24/06/2023 | 158.594 | -21.0313 | 26.7 | -0.4 [-0.6,-0.3] | 23 |
| New Caledonia | Ilot Loop | CB3 | 25/06/2023 | 158.4083483 | -19.9285188 | 27.3 | -0.3 [-0.4,-0.1] | 24 |
| New Caledonia | Chesterfields SO | CB4 | 26/06/2023 | 158.478 | -19.905 | 27.3 | 1.1 [0.4,2.5] | 17 |
| New Caledonia | Chesterfields NO | CB5 | 27/06/2023 | 158.433288 | -19.03805 | 27.7 | 1.8 [-0.1,4.1] | 18 |
| New Caledonia | Chesterfields NE | CB6 | 28/06/2023 | 158.884708 | -18.960638 | 27.6 | 2.5 [0.6,5] | 22 |
| New Caledonia | Recif du Prony | NC1 | 10/08/2023 | 166.3273667 | -22.26615 | 26.6 | 1.1 [-0.2,4.1] | 25 |
| New Caledonia | Recif Mara | NC2 | 22/08/2023 | 165.72805 | -21.830483 | 26.9 | 3 [0.9,5.6] | 24 |
| New Caledonia | Recif Snark | NC3 | 01/09/2023 | 166.42516 | -22.44361 | 26.6 | 0.4 [0,1.4] | 21 |
| New Caledonia | Ile Verte | NC4 | 06/09/2023 | 165.47735 | -21.68475 | 27 | -0.4 [-0.5,-0.1] | 25 |
| New Caledonia | Ilot Contrariete | NC5 | 13/09/2023 | 165.05492 | -21.46439 | 27.1 | 0.9 [0.4,1.7] | 25 |
| New Caledonia | Ilot Deverd | NC6 | 19/09/2023 | 164.3237 | -20.766433 | 27.5 | 0.3 [-0.4,1.9] | 25 |
| New Caledonia | Plateau de Koniene | NC7 | 20/09/2023 | 164.7209 | -21.157383 | 27.3 | 0.2 [-0.2,1.9] | 25 |
| New Caledonia | Pointe de Poum | NC8 | 28/09/2023 | 163.99965 | -20.24595 | 27.8 | -0.2 [-0.5,1.3] | 25 |

**Table S2.** Pre-variant filtering steps summary. Same filtering parameters were used to process raw sequence reads from GBR (Batch 1) and New Caledonia (Batch 2) samples.

| Filtering step | Statistic | Batch 1 (GBR samples) | Batch 2 (NC samples) |
|---|---|---|---|
| **Samples collection, selection and sequencing** | N initial samples collected | 831 | 323 |
| | N initial samples sequenced | 830 | 314 |
| | N technical replicates | 2 | 23 |
| | N other species samples included as outgroups | 36 | 0 |
| **Sequence QC** | **Total number of samples successfully sequenced** | **868** | **337** |
| | Total number of reads prior filtering | $31.7 \times 10^9$ | $14.2 \times 10^9$ |
| | Total number of reads after quality filtering | $29.8 \times 10^9$ | $12.3 \times 10^9$ |
| **Mapped reads filtering** | Number of reads after filtering for PCR and optical duplicates | $25.3 \times 10^9$ | $10.5 \times 10^9$ |
| | Number of reads that mapped | $24.3 \times 10^9$ | $97.3 \times 10^8$ |
| | Number of reads that primarily mapped | $23.1 \times 10^9$ | $92.4 \times 10^8$ |
| | Number of reads after filtering improperly paired reads | $20.5 \times 10^9$ | $80.7 \times 10^8$ |
| | **Number of samples after filtering out <80% and <10M reads mapping** | **859 (including 821 initially collected colonies)** | **309 (including 286 initially collected colonies)** |
| **Taxa mis-identification removal** | Number of samples identified as distinct from target taxa | 4 | 23 |
| | **Total number of samples used for SNP and variant calling (after removal of other taxa)** | **819** | **286** |

**Table S3.** Post-variant filtering steps summary. For each step, the number of retained samples and single nucleotide polymorphisms (SNPs) is reported.

| Step | N samples | N SNPs |
|---|---|---|
| Joint genotyping | 1,105 | 92,885,427 |
| GATK VariantFiltration (QD, FS, MQ, SOR, MQRankSum | 1,105 | 47,438,630 |

| | | |
|---|---|---|
| Filtration of individuals with high missingness (>85%) | 1,088 | 47,438,630 |
| Preliminary filtration (mac>1, site missingness <20%, bi-allelic SNPs…) | 1,088 | 339,720 |
| Removal of technical replicates and individuals with >0.2 relatedness | 999 | 339,720 |

**Table S4.** SNPs and genotypes filtering parameters used to conduct the main analyses. Preliminary analyses including variable missingness rates between 5 and 50% showed consistent results with the final threshold used to report the results (20%).

| **Filtering step** | **Population structure (Dataset 1)** | **ML tree (Dataset 1)** | **Population statistics computation (Dataset 2)** | **Demographic modeling (Dataset 3)** | **Isolation by distance/$N_e$ (Dataset 4)** |
|---|---|---|---|---|---|
| Min GQ | 20 | 20 | 20 | 20 | 20 |
| Min Genotype coverage | 5 | 5 | 5 | 5 | 5 |
| Max Genotype coverage | 15 | 15 | 50 | 15 | 500 |
| Scope | Study-wide | Study-wide | Between population pairs and within pops | Between population pairs | Within & Across population |
| Number of samples | 999 | 280 (subset of 70 samples per pop) | 20 x 10 (subset of 5 samples per pop x 10 replicates) | 120 (subset of 30 samples per pop) | 999 |
| Max missing per locus | 20% | 20% | <20% | <20% within each population | 10% |
| MAF | MAF>0.05 | MAF>0.05 | | | |
| MAC | MAC>1 | MAC>1 | MAC>1 | MAC>3 | MAC>3 |
| HWE p-values | | | p-value>0.0001 | p-value>0.0001 | |
| Physical pruning | | | | | 1 SNP per 500bp |
| LD | LD<0.2 | LD<0.2 | | | |
| Number of SNPs | 27,175 | 27,175 | ~2.2M SNPs + invariant sites | 37,841-61,902 | 2,011 |

**Table S5.** Environmental predictors considered for Symbiodiniaceae community distance-based redundancy analysis and host structure redundancy analysis. Predictors in bold are those retained after controlling for multicollinearity (VIF<10) and used in forward stepwise selection.

| Variable | Metric | Source | Spatial resolution | Time resolution | Period | Description |
|---|---|---|---|---|---|---|
| Temperature | DHW_avg | Coralwatch | 5km | daily | 1985-2024 | Average monthly maximum DHW |
| | ***SST_AR*** | Coralwatch | 5km | daily | 1985-2024 | SST Annual range |
| | SSTmean_change | Coralwatch | 5km | daily | 1985-2024 | Nonparametric effect size between the distribution of monthly temperature between decade 1985-1995 and 2013-2023 |
| | ***MMM*** | Coralwatch | 5km | daily | 1985-2024 | Average temperature of the warmer calendar month (1985-1990+1993 climatology) |
| | ***DHW_freq_sup4*** | Coralwatch | 5km | daily | 1985-2024 | Average number of days above 4 DHW per year |
| | **DHW_max** | Coralwatch | 5km | daily | 1985-2024 | Maximum DHW |
| | SST_average | Coralwatch | 5km | daily | 1985-2024 | Average SST |
| UV radiation | UVR_median | ERA5 | 0.25° | hourly | 2002-2024 | Median UV radiation (W.m $^{-2}$) |
| Irradiance | **CF_median** | ERA5 | 0.25° | hourly | 2002-2024 | Median cloud fraction |
| Water clarity | **Kd490_median** | Globcolour | 1/24° | daily | 2002-2024 | Median attenuation of irradiance at 490nm, proxy for turbidity |
| Chlorophyll | Chla_median | Globcolour | 1/24° | daily | 2002-2024 | Median concentration of chlorophyll a = proxy for nutrient load (mg. m$^{-3}$) |
| Spatial | **Km_to_coastline** | GADM 4.1 | | | | Shortest haversine distance to the shore |
| | Lat | | | | | Latitude hexadecimal |

**Supplementary Figures**

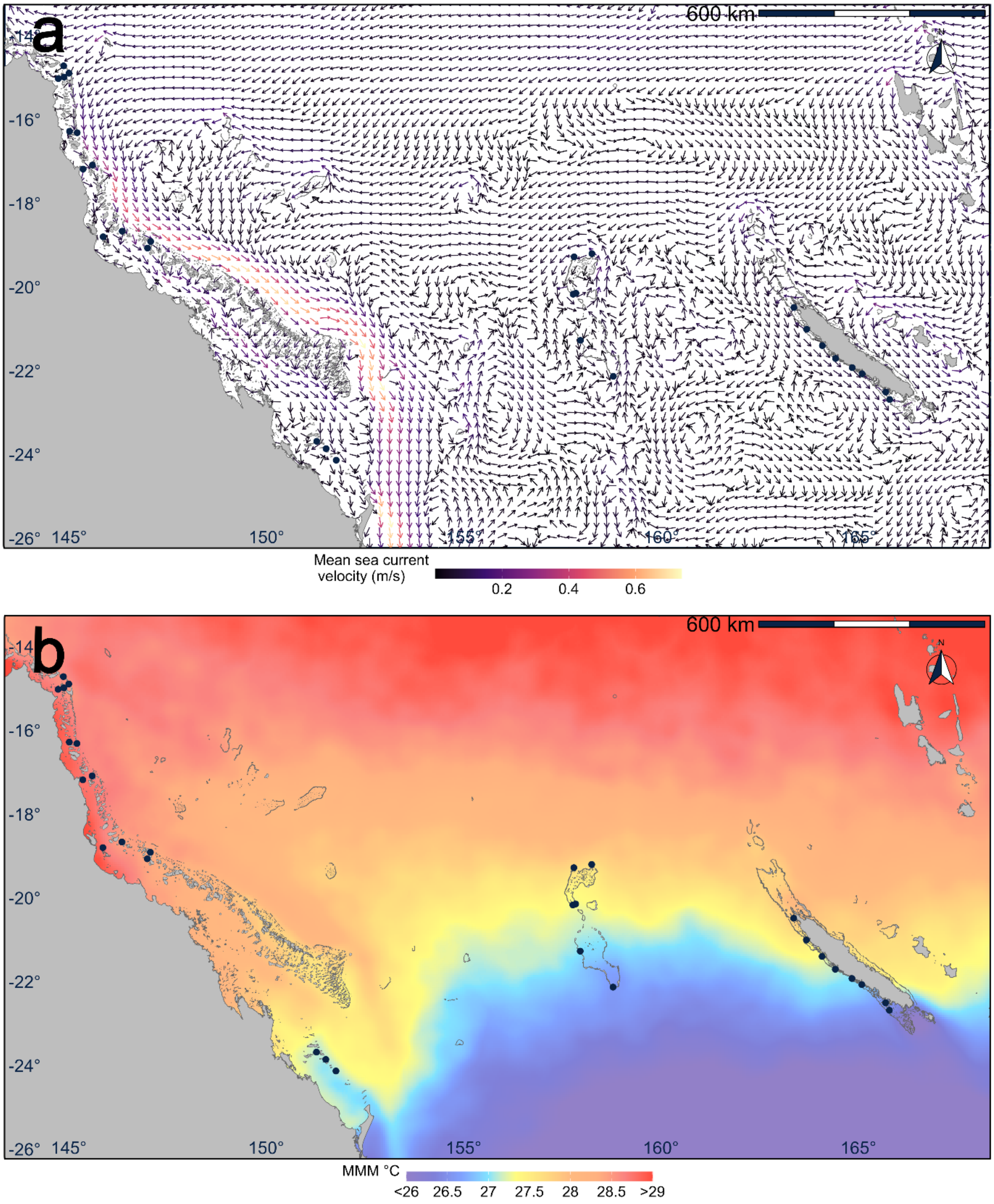


**Figure S1.** Environmental conditions across the Coral Sea. a) Mean surface sea currents ($m.s^{-2}$) averaged over known spawning months for *A.spathulata* : October, November, December for the period 1993-2021 obtained from Copernicus GLOBAL_MULTIYEAR_PHY_001_030 product. b) Maximum Monthly Mean (MMM, °C) obtained from NOAA CoralWatch v3.1. Black dots indicate *Acropora spathulata* sampling sites.

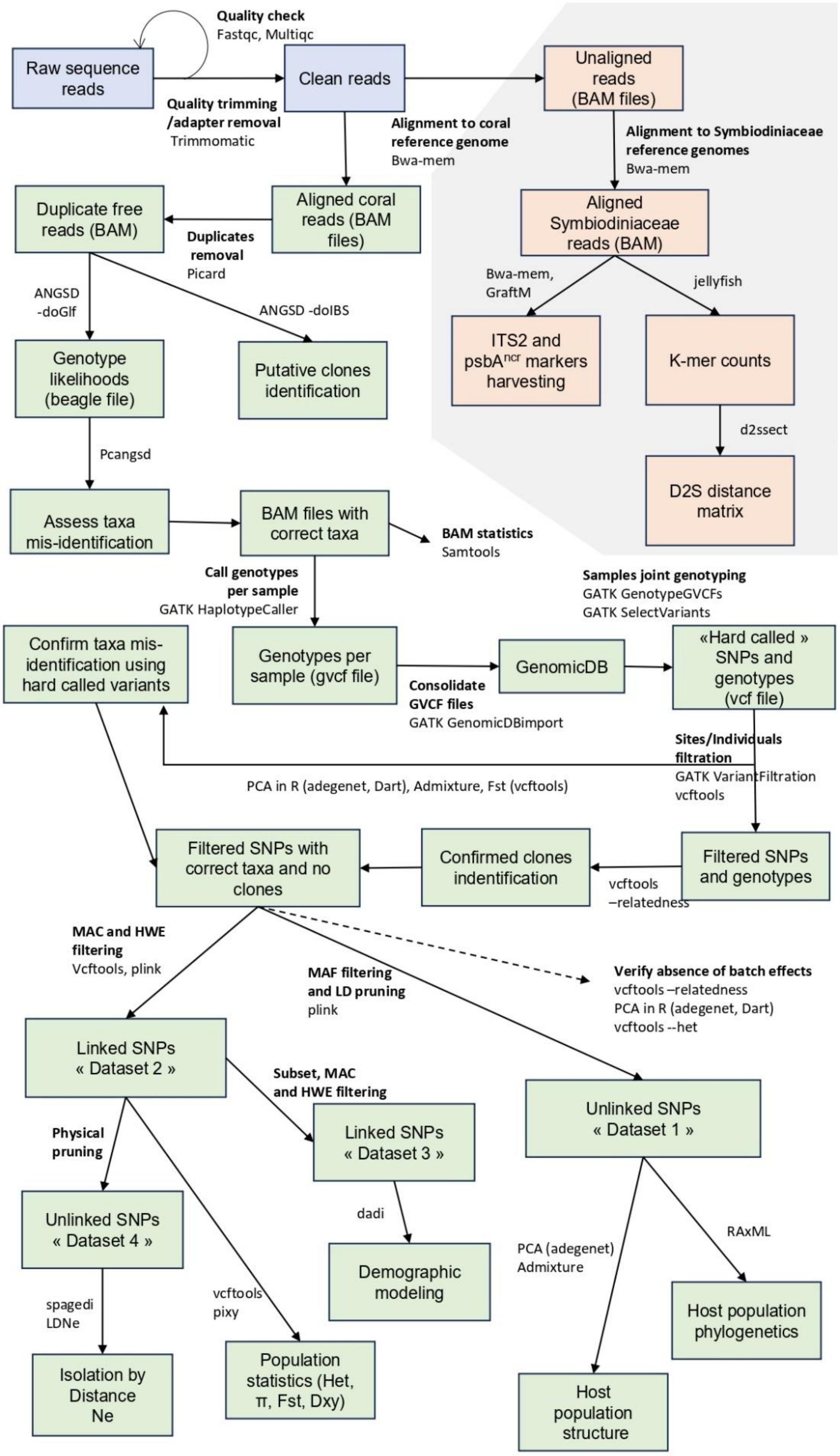


**Figure S2.** Summary of bioinformatic analyses workflow used to assess the population structure of coral hosts and their symbiont communities across the Coral Sea. Green boxes depict data processing steps for the host and red boxes depict data processing steps for the symbionts.

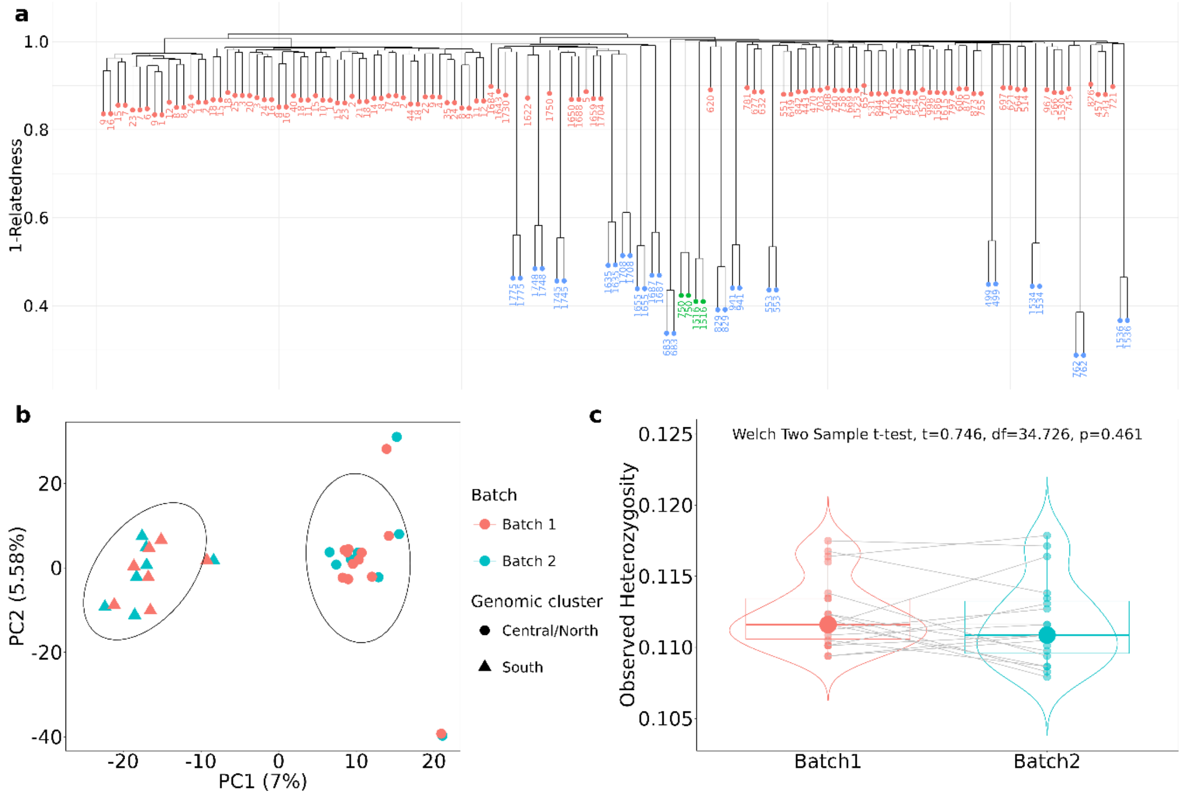


**Figure S3.** Assessment of potential batch effects using technical replicates extracted and sequenced within each batch. (a) Relatedness and (b) Principal component analyses were performed using 130,344 polymorphic sites (20% missingness). (c) Individual autosomal heterozygosity was computed from 2,715 polymorphic and monomorphic sites with 0% missing data, following recommendations of Schmidt et al. (2021).

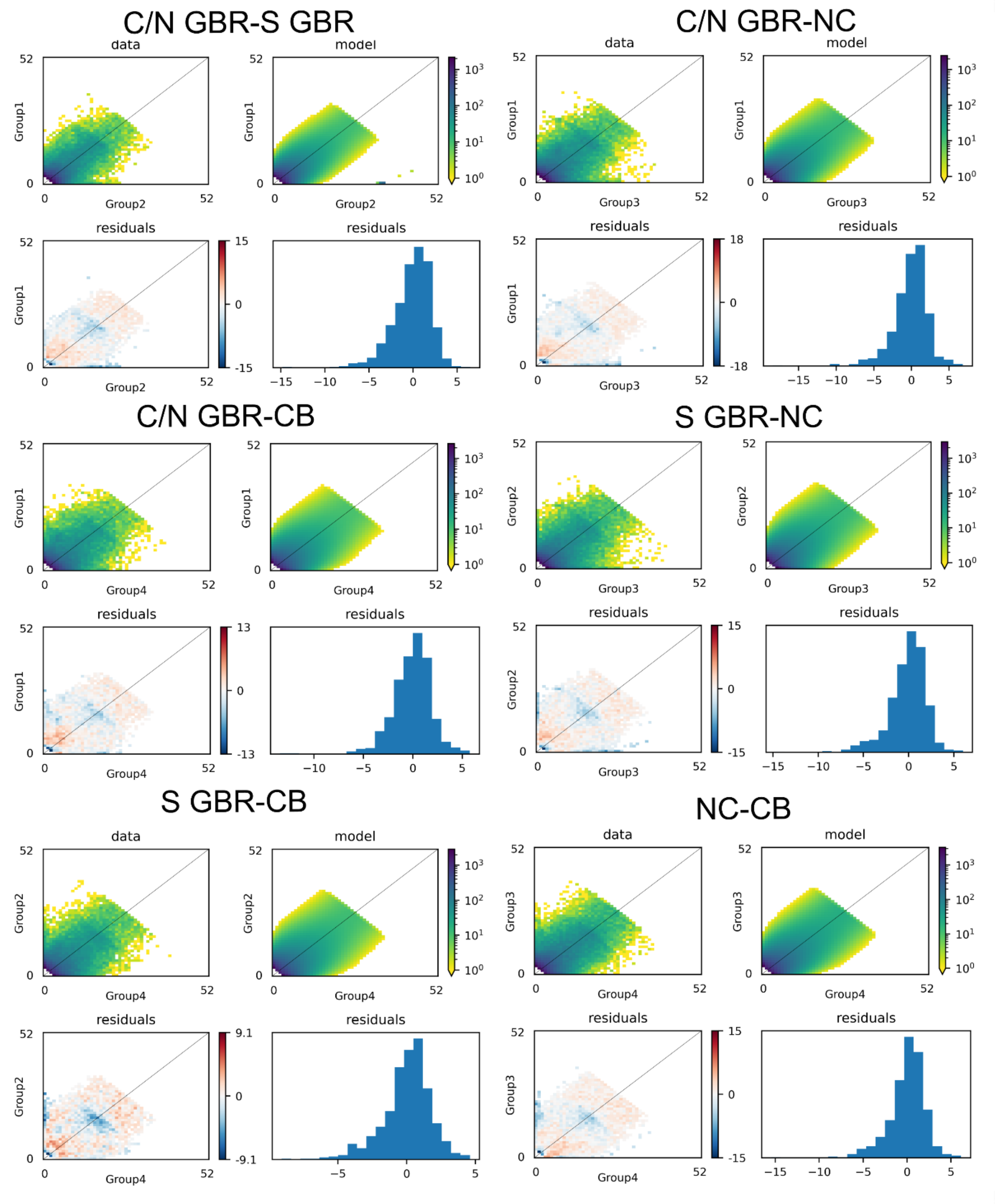


**Figure S4.** Observed and Modeled joint Allelic frequency spectrum (JAFS) for each pair of genomic populations (11 individuals per population) using demographic modeling analyses in Dadi. For each pair of genomic populations panels show observed JAFS (top left), best modeled JAFS (top right), and residuals distribution (bottom plots). Low frequency alleles are masked to account for genotyping errors.

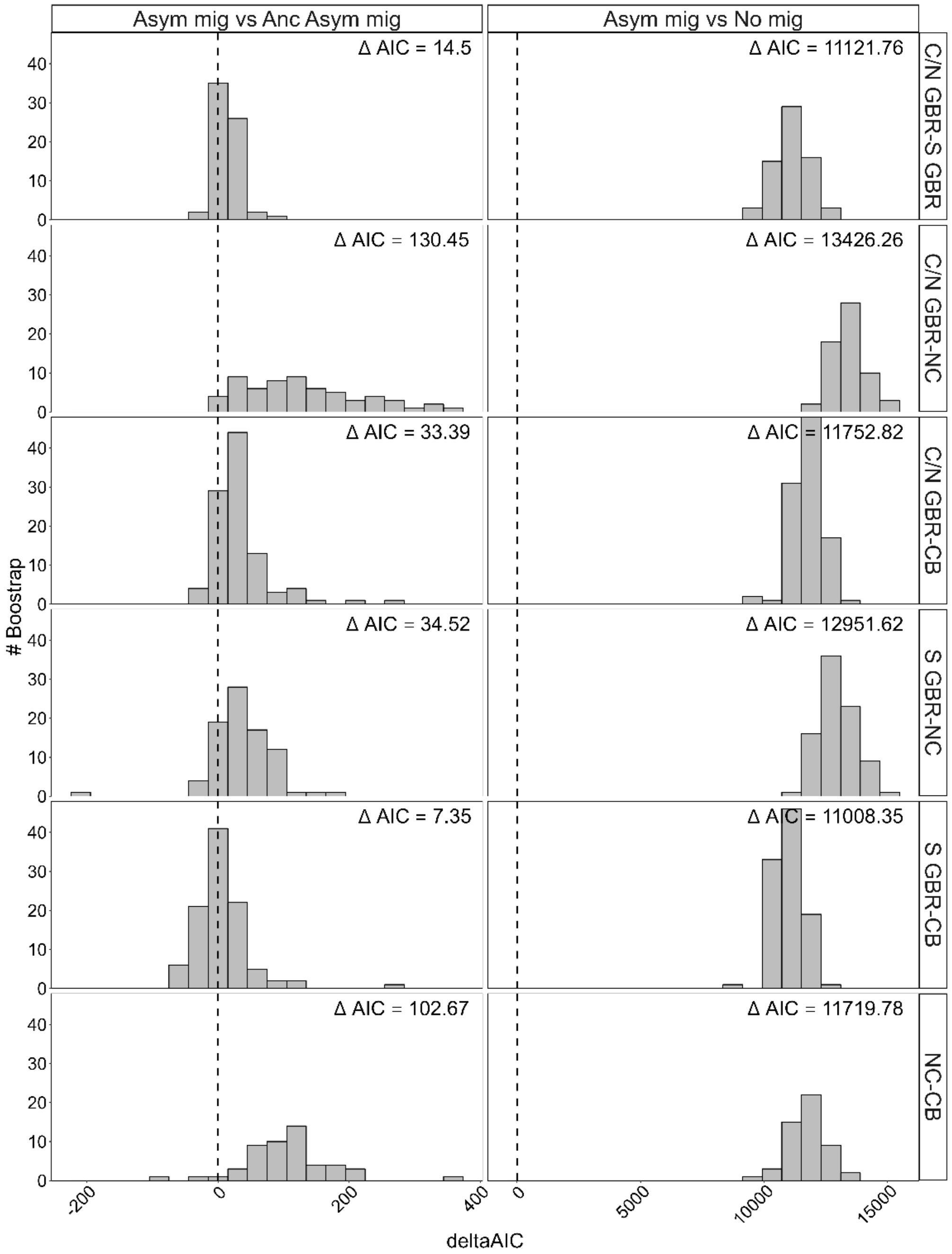


**Figure S5.** Bootstrap analysis of *A. spathulata* populations divergence models. Histograms of delta-AIC values for bootstrap replicates comparing a model of divergence with asymmetrical gene flow to a model of divergence with ancestral gene flow (left, gene flow stopped at some time T in the past) and divergence in the absence of gene flow (right). The mean delta-AIC across 100 bootstraps is indicated for each population and model comparison. Positive values indicate higher support for a model of divergence with asymmetrical gene flow.

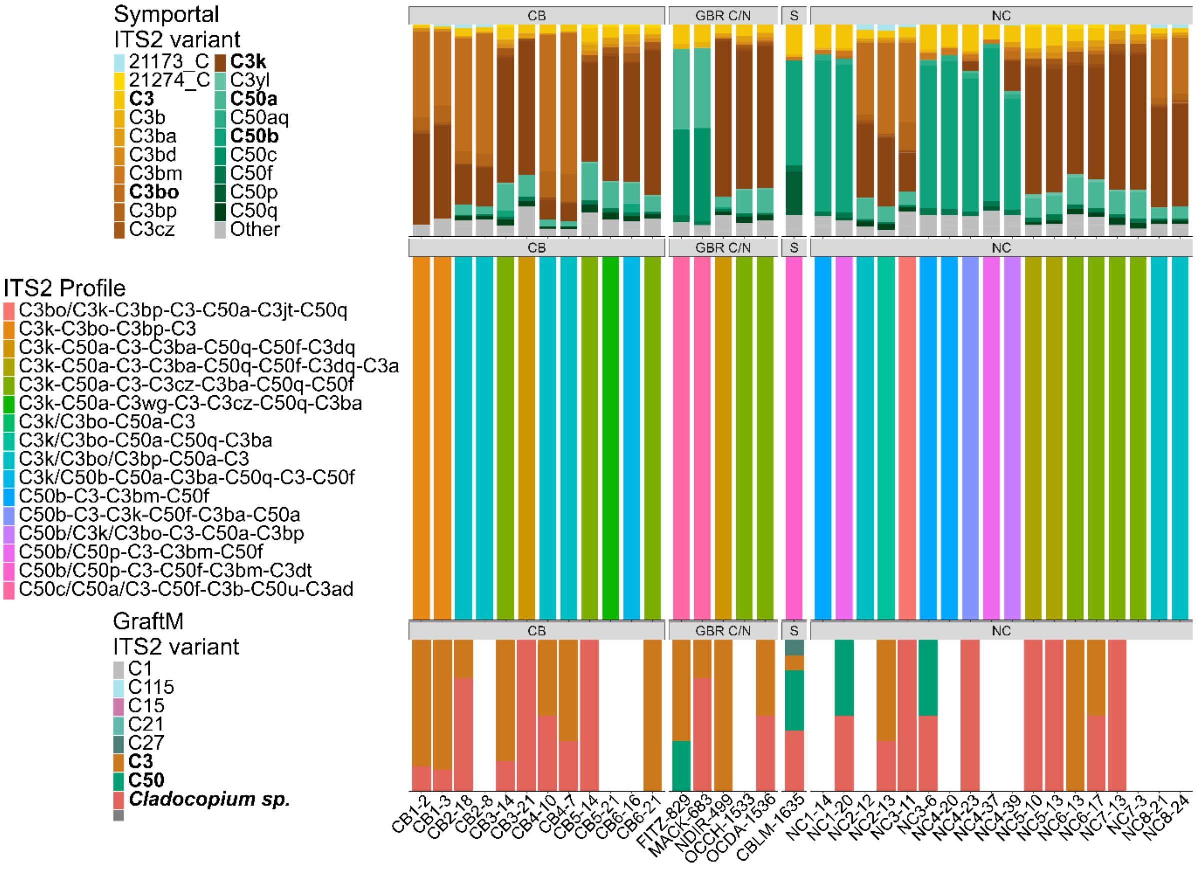


**Figure S6.** Comparison of ITS2 sequence variants recovered using graftM in hologenome data with conventional amplicon sequencing analyzed in Symportal. Despite the low number of reads recovered by graftM both identified similar dominant ITS2 subclade in samples as C3 or C50.

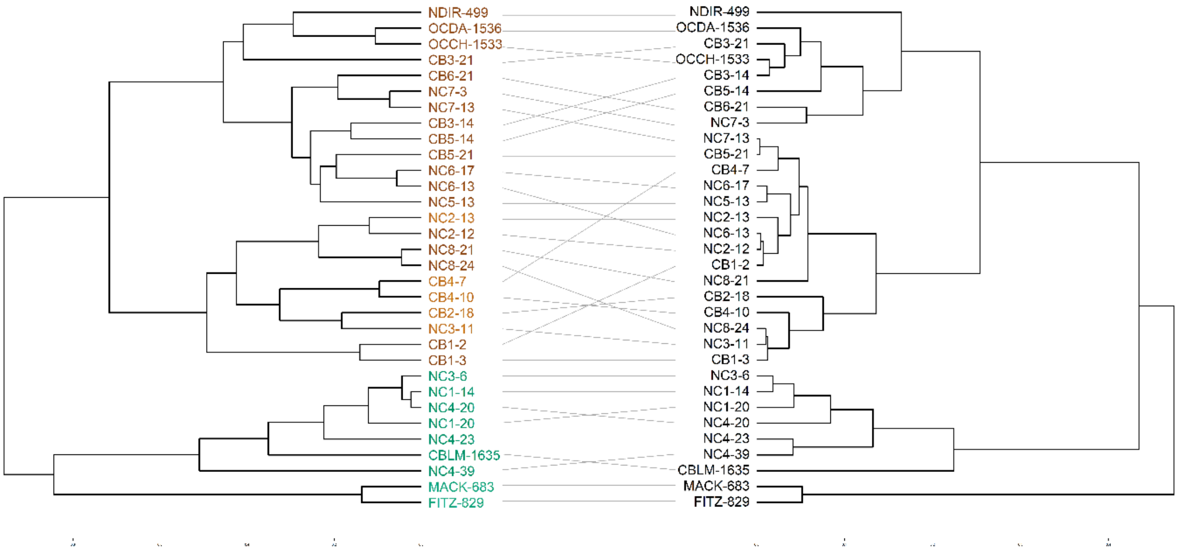


**Figure S7.** Comparison of samples hierarchical clustering using Unifrac distance based on ITS2 ‘defining intragenomic variants’ (DIV, left) and D2S distance based on k-mer profiles (right). Leaves in the left dendrogram are colored according to the majority ITS2 sequence found in the sample (C3k, C3k/C3bo, C50c or C50b).

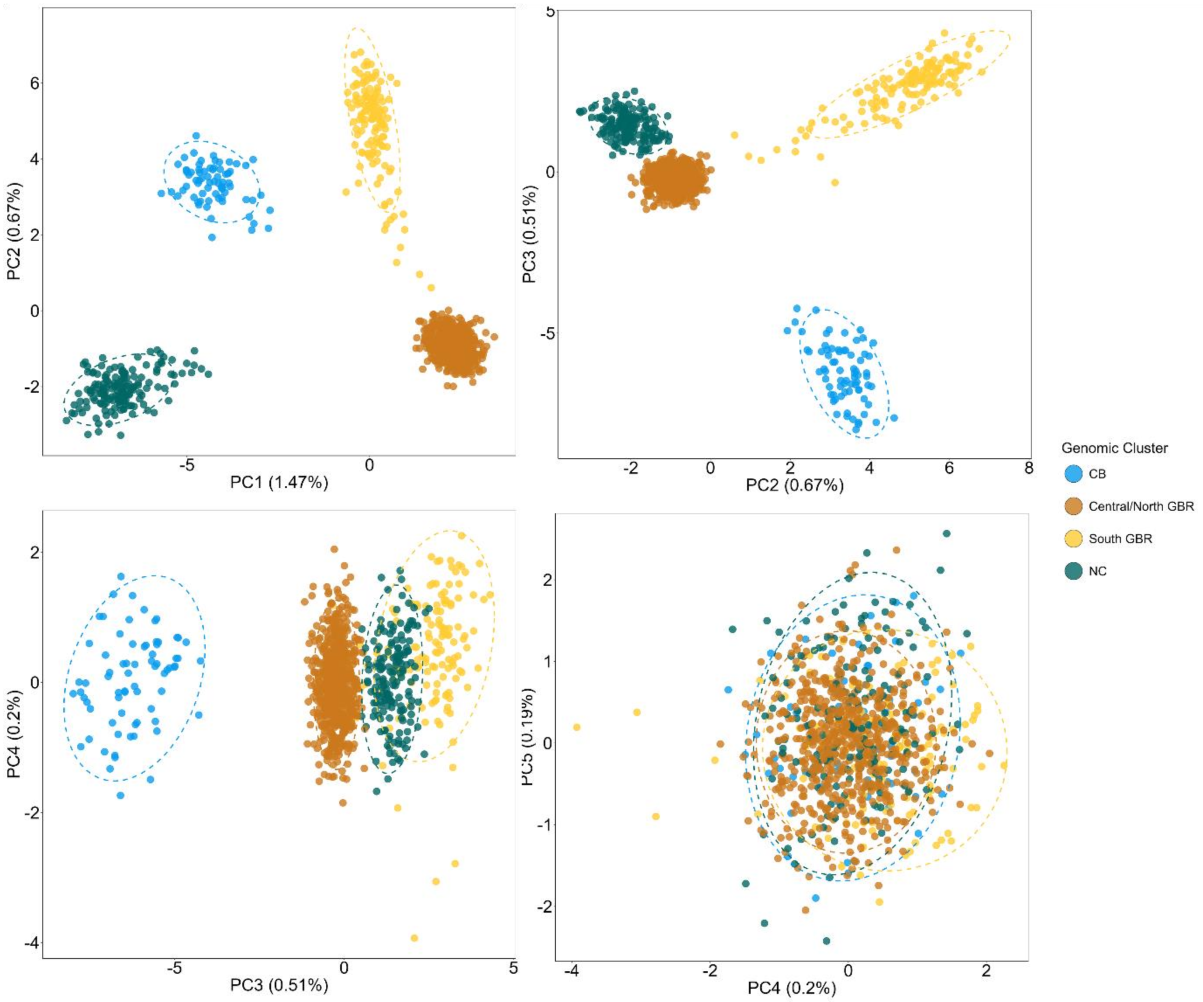


**Figure S8.** Principal component analyses on 27,175 high quality 'hard called' SNPs from 999 colonies collected across the Coral Sea. Each point represents an individual colored according to its geographic region (CB = Chesterfields-Bellona atolls, NC = New Caledonia, GBR = Great Barrier Reef). The first 5 principal components are shown with number in parenthesis representing the % of total variance explained by that component.

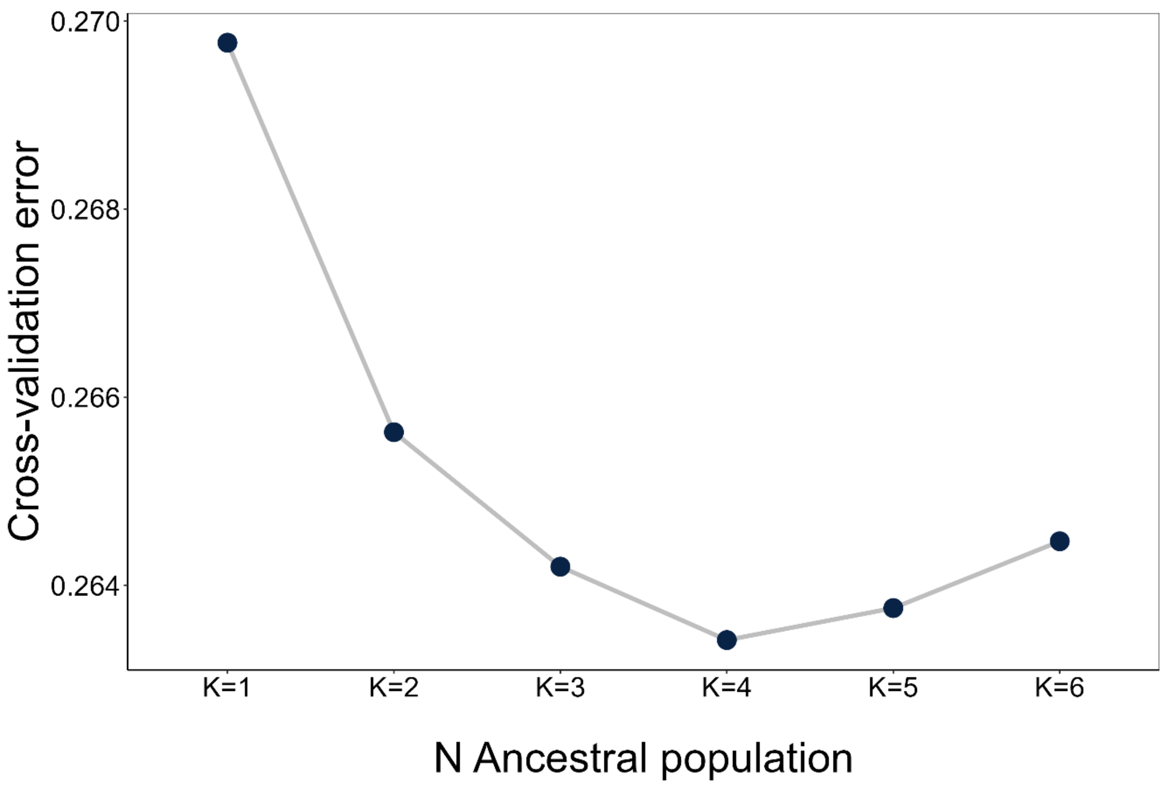


**Figure S9.** ADMIXTURE cross-validation error for different numbers of ancestral population. The optimal choice (K=4) showing the lowest error was used to conduct the final analyses.

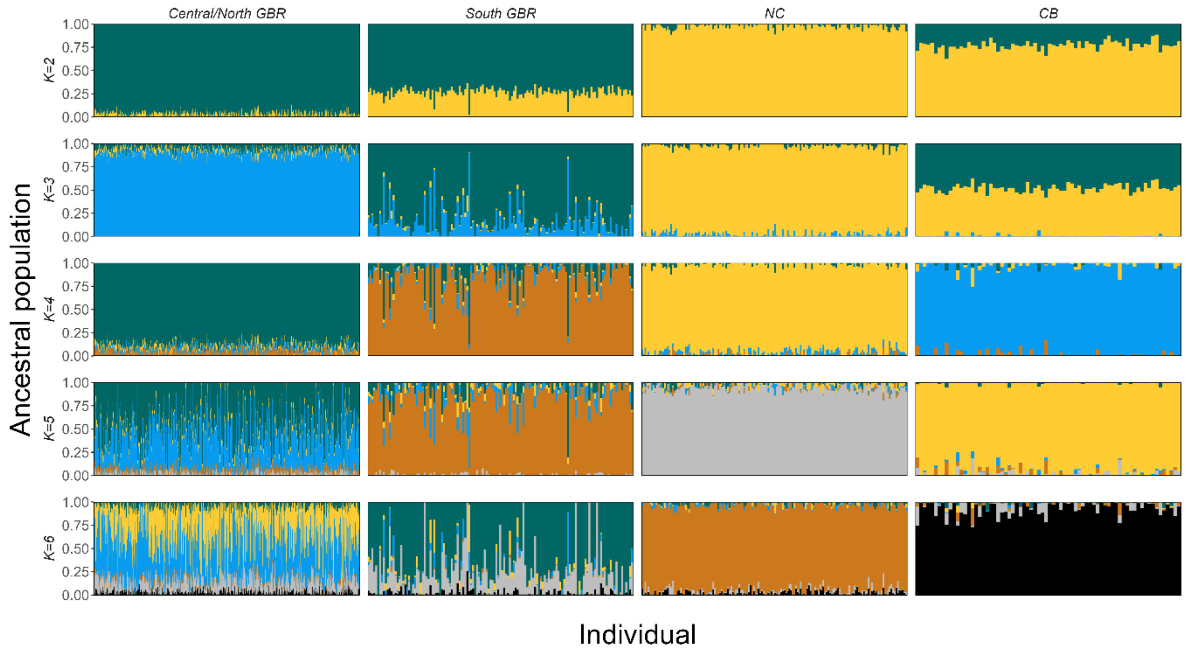


**Figure S10.** Ancestral population proportion for each of 999 colonies across the coral sea using ADMIXTURE analysis on 27,175 ‘hard called’ SNPs and K=2–6 ancestral population.

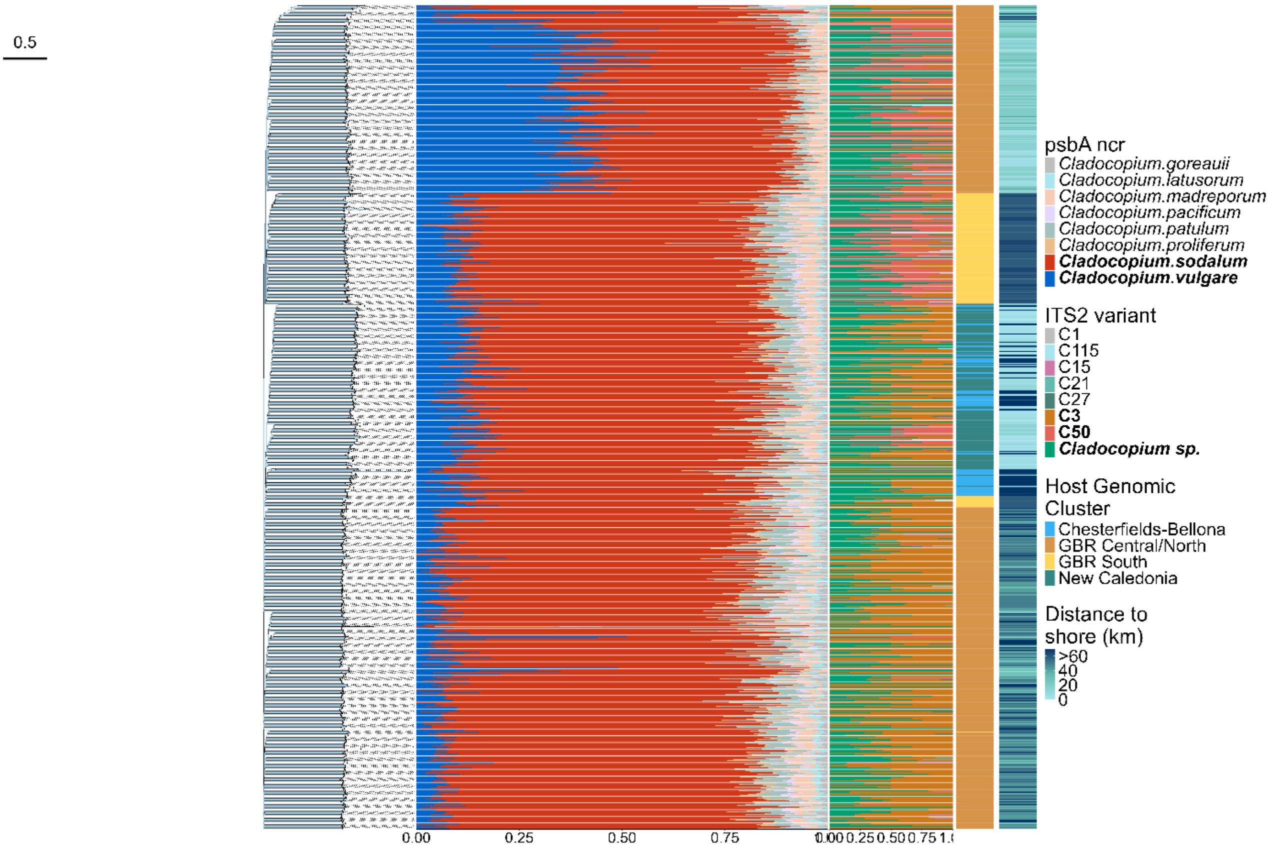


**Figure S11.** Summary of marker-based and alignment-free approaches used to characterize Symbiodiniaceae genetic diversity. A neighbour-joining tree was obtained based on D2S distance computed from k-mer pofiles (left). psbA$^{ncr}$ sequences were aligned to a custom reference database and the relative abundance of uniquely mapping sequences is shown (center). ITS2 sequences were recovered in the samples using graftM and classified using symportal reference database (right). Colored tiles show assignment to host genomic populations and cross-shore gradients.

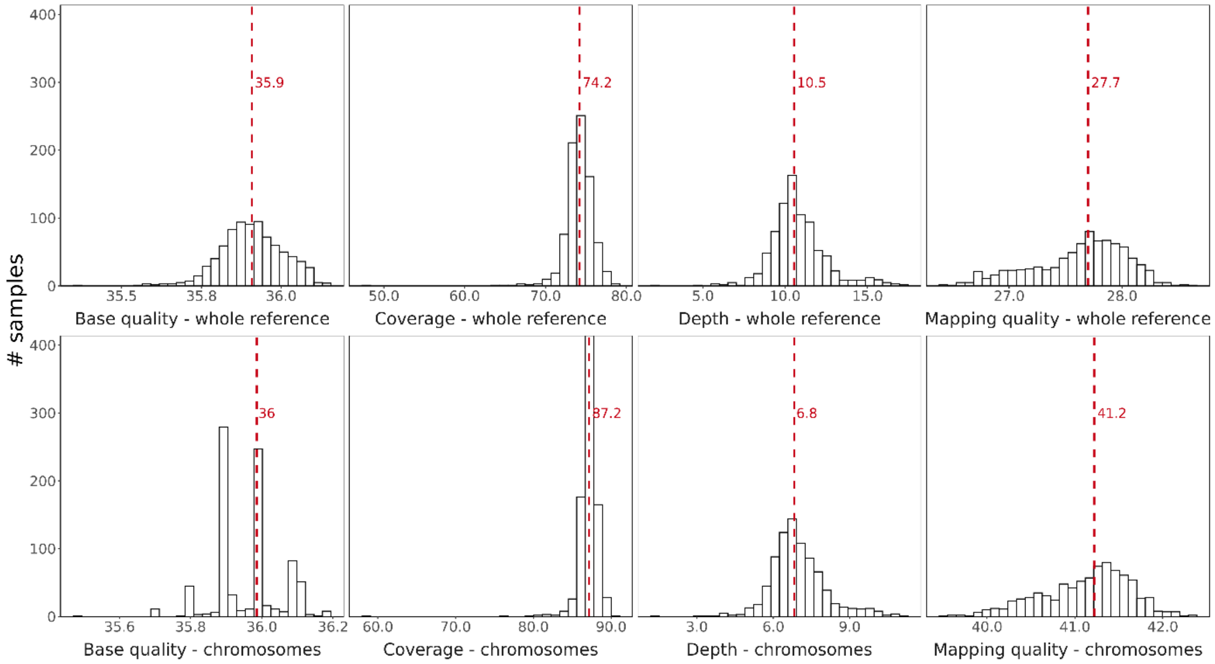

**Figure S12.** Summary of BAM files statistics for GBR samples (Batch1). Statistics are reported for the whole reference (chromosome + contigs) and for chromosomes only. Only reads mapping to the chromosomes were used in further analyses.

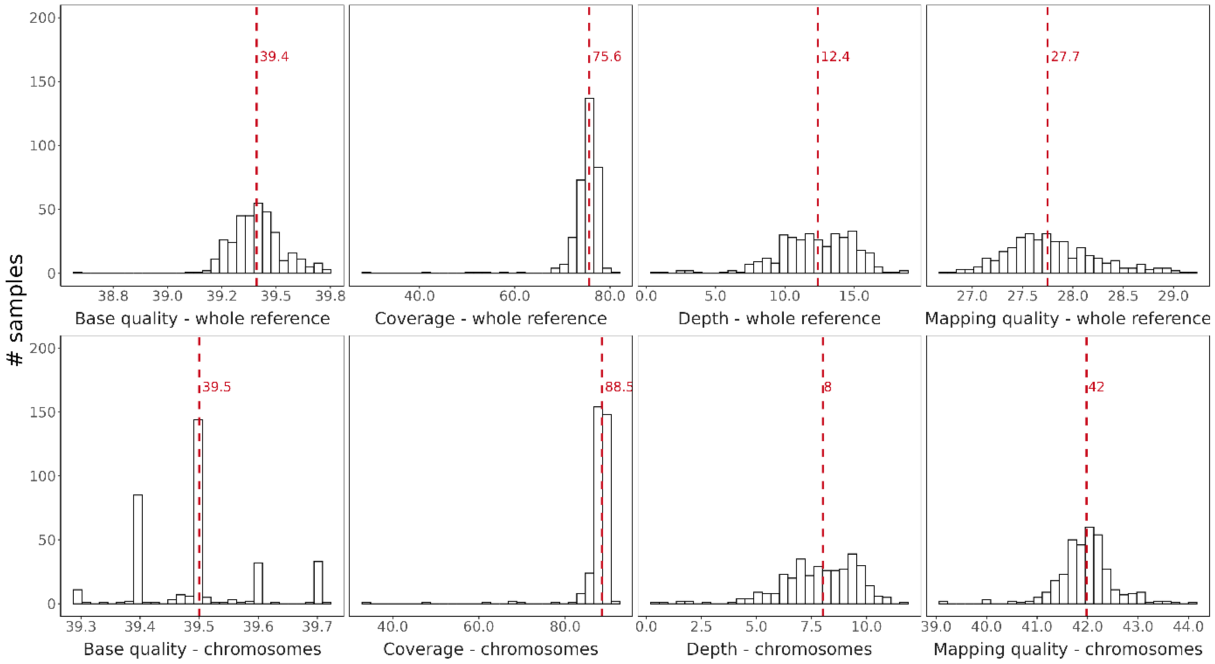


**Figure S13.** Summary of bam files statistics for NC samples (Batch2). Statistics are reported for the whole reference (chromosome + contigs) and for chromosomes only. Only reads mapping to the chromosomes were used in further analyses.

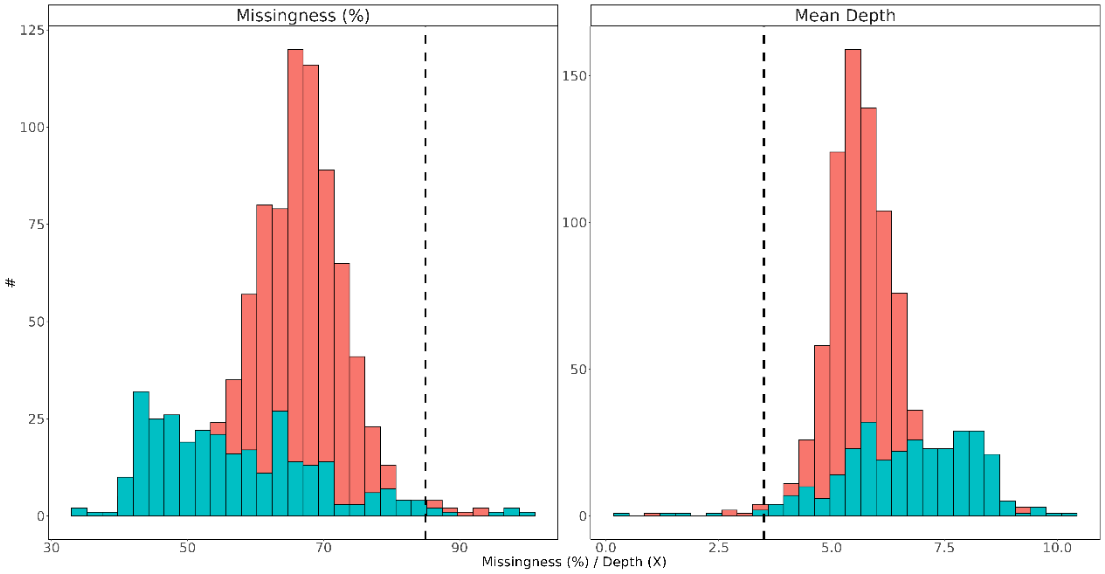


**Figure S14.** Distribution of individual missingness and depth across 1,105 individuals and 47,438,630 SNPs for Batch 1(pink) and Batch 2 (blue). The vertical dashed lines represent the thresholds used to remove individuals with high missingness and/or low average depth resulting in 1,088 high quality individuals (17 individuals removed).

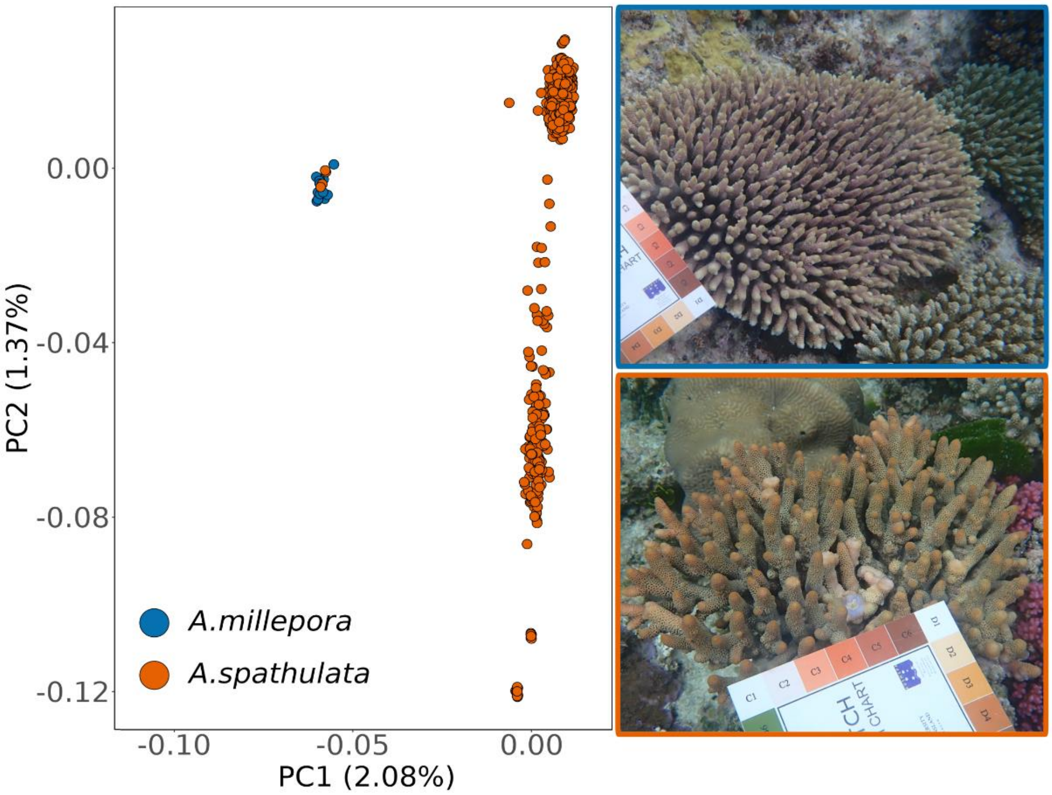


**Figure S15.** Taxa mis-identification within the GBR dataset. Four samples were identified as *A.millepora* and discarded from following analyses. Pictures on the right show representative examples of *A.millepora* (blue) and *A.spathulata* (orange) colonies at Heron Island.

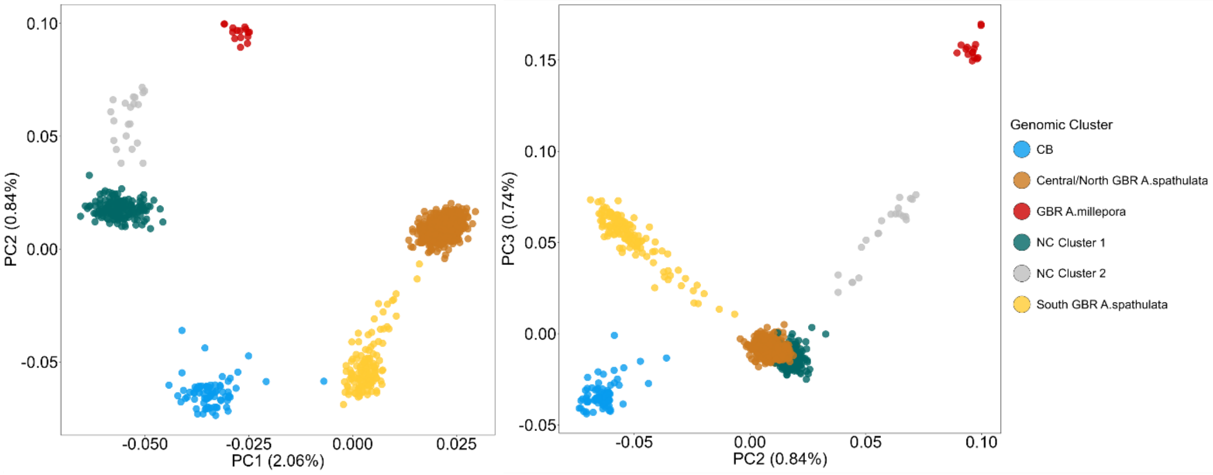


**Figure S16.** Principal component analysis of the global dataset alongside *A.millepora* samples from the Great Barrier Reef using genotype likelihood (1,041 individuals and 1,474,993 polymorphic sites).

The putative undersampled Cluster 2 from New Caledonia and *A.millepora* outgroup were discarded from further analyses.

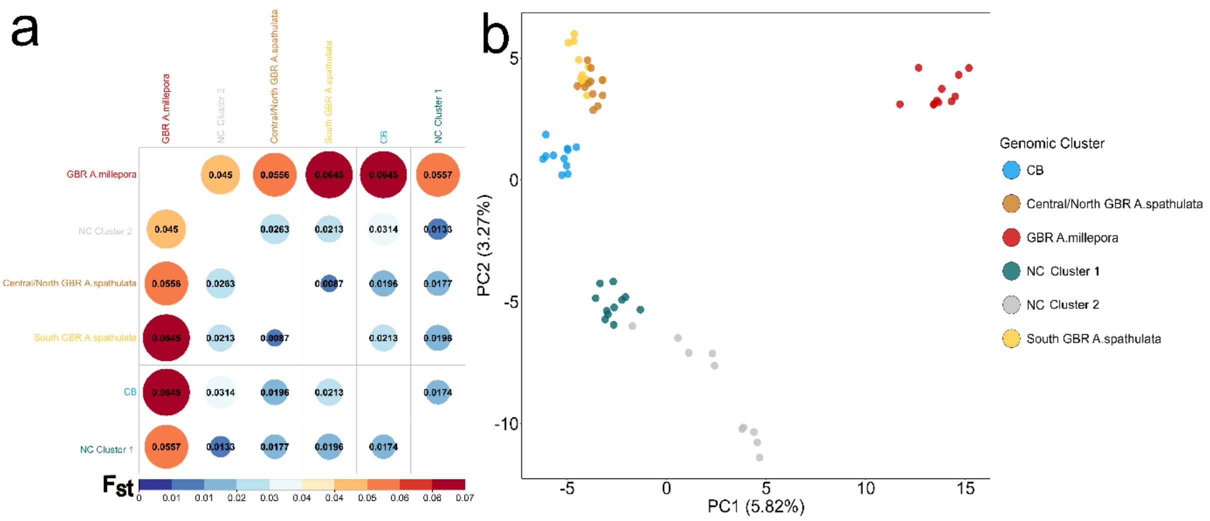


**Figure S17.** Genetic differentiation between groups using hard-called variants. Pairwise fixation index ($F_{st}$) computation (a) and Principal Component Analysis (b) were performed on 7,051 SNPs and a reduced dataset of 66 colonies (11 colonies per group). The putative undersampled Cluster 2 from New Caledonia and *A.millepora* outgroup were discarded from further analyses.

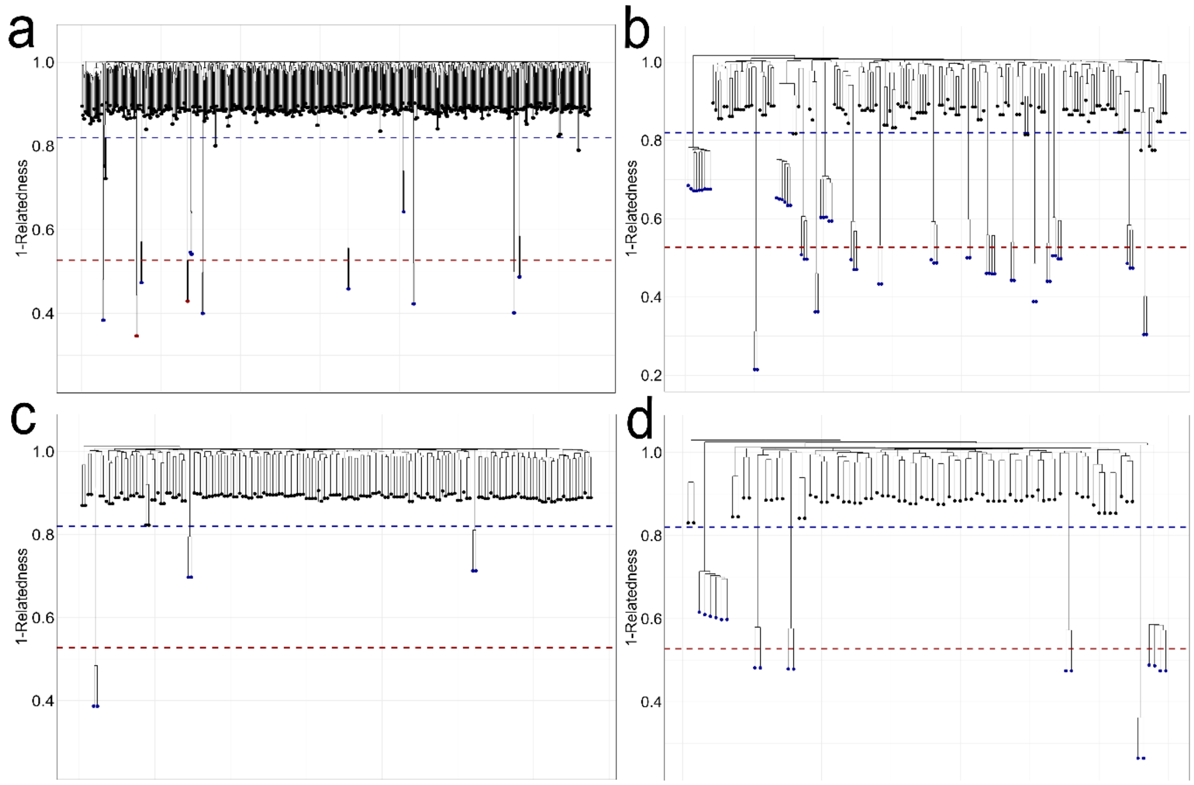


**Figure S18.** Genomic relatedness among individuals for Central/North GBR (a), South GBR (b), NC (c) and CB (d) genomic populations. The red line shows the minimum relatedness among technical replicates and the blue line shows the conservative threshold (r>0.2) used to remove clones and related individuals (blue leaves) from the dataset for further analyses.

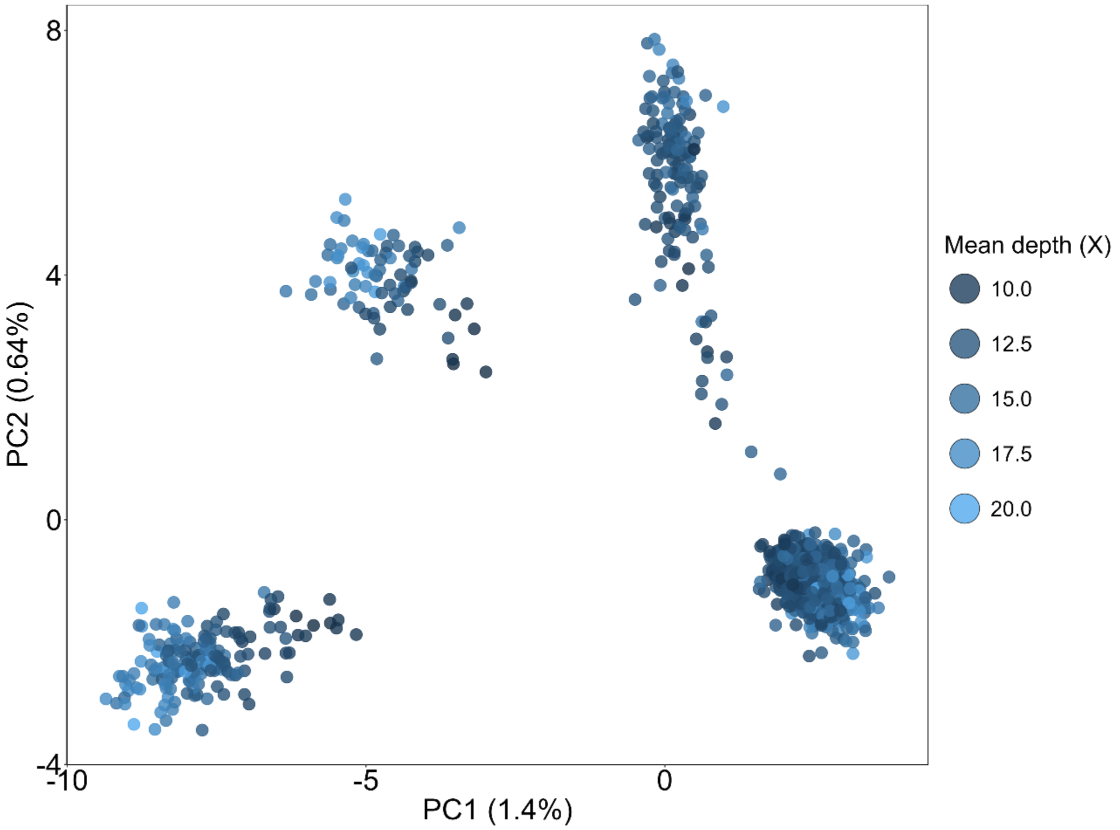


**Figure S19.** Principal Component Analysis on 27,175 high quality 'hard called' SNPs from 999 colonies collected across the Coral Sea. Each point represents an individual and is colored by its average sequencing depth across all loci to control for potential bias in population structure. The first two principal components explain 1.4% and 0.64% of the total variance respectively.

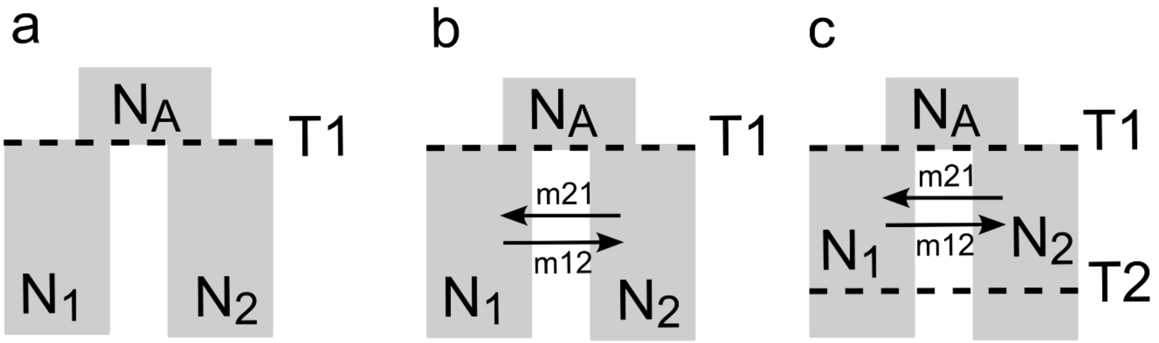

**Figure S20.** Models considered in demographic modeling analyses: a) divergence in isolation, b) divergence with asymmetrical homogeneous gene flow, c) ancestral asymmetrical migration. Preliminary models of no divergence were also tested.

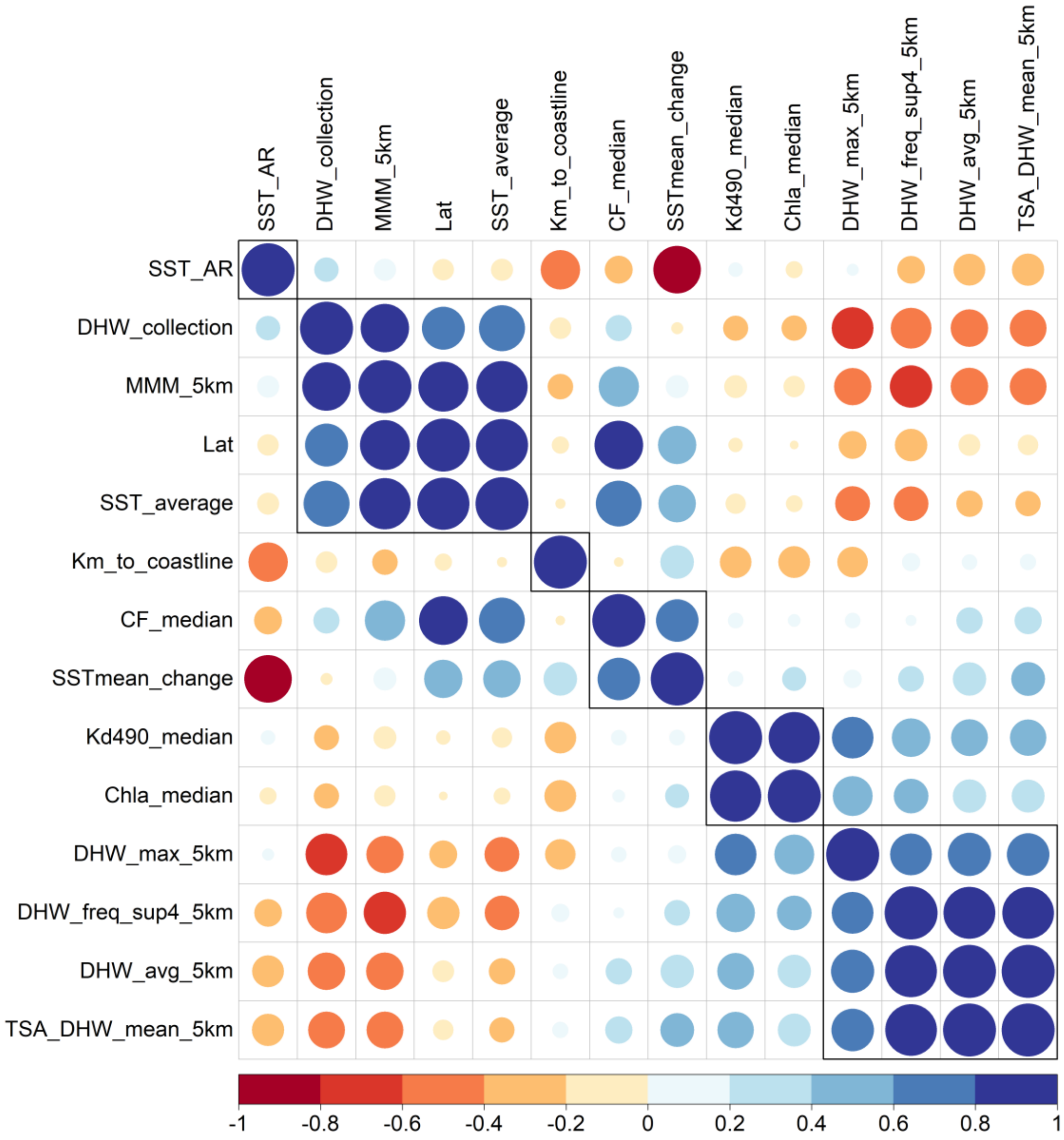


**Figure S21.** Pairwise Pearson's correlation coefficients between the 14 quantitative environmental predictors considered for the redundancy analyses. Only 7 predictors with a variance inflation factor < 10 were used for stepwise forward selection in the final model.
